# Beyond single-trait analysis: ancient genomes reveal coordinated evolution of dietary endophenotypes in Holocene Europe

**DOI:** 10.64898/2026.10.27.762183

**Authors:** Carla Casanova-Suárez, Karen Hardy, Oscar Lao

## Abstract

Diet has been a major selective force throughout human evolution, yet dietary adaptation is often studied through individual loci rather than as interconnected systems of traits. Here, we integrated genome-wide association data for 58 diet-related phenotypes with genomes from 938 ancient Europeans spanning the last 10,000 years to reconstruct the evolution of dietary endophenotypes. We identified four major systems encompassing adiposity and metabolism, body composition, fatty-acid metabolism and food-related behaviours. Multiple endophenotypes showed significant temporal shifts during the Holocene departing from the expected under neutrality, including coordinated decreases in genetic architectures associated with adiposity, insulin regulation and lipid metabolism, alongside increases in traits related to docosahexaenoic acid metabolism. Moreover, these patterns showed marked geographic heterogeneity across Europe. Our results indicate that dietary adaptation proceeded through coordinated evolutionary changes across interconnected physiological systems, providing a systems-level perspective on how cultural and subsistence transitions reshaped human biology.

## Introduction

Diet has been a major ecological pressure in human evolution, shaping how human populations obtain, process and metabolize energy across changing environments. In particular, the Holocene transition from foraging to farming and pastoralism transformed human nutritional landscapes through changes in plant and animal resource use, cereal and starch intake, dairying, lipid sources, food storage and subsistence scheduling^1–3^. Specifically, this transition is among the most frequently acknowledged examples of human adaptation and gene-culture coevolution^4^. Some of the clearest examples of strong recent selective pressures reported in humans involve single diet-related loci, including lactase persistence at *LCT*, copy-number variation at *AMY1* linked to starch digestion, or variation near FADS genes involved in fatty-acid metabolism^2,3^. However, dietary adaptation has often been investigated through a limited set of predefined dietary variables or broad subsistence categories rather than as a multidimensional phenotype composed of interacting nutritional, ecological, cultural, and behavioural components^5^.

Human dietary behaviour is not a single phenotype but encompasses multiple interconnected biological and behavioural traits^5–9^. Genome-wide association studies (GWAS) have shown that most human phenotypes –including diet-related– are complex shaped by the combined effects of many genetic variants, environmental factors, and interactions among biological systems^10,11^. The genetic architecture of complex traits is typically highly polygenic, and that genetic variants frequently influence multiple traits, generating extensive pleiotropy and genetic correlations among phenotypes^10–12^. Consequently, evolutionary histories of diet-related traits are non-independent and share genetic variants, whose alleles can have concordant or opposing effects on temporal changes across phenotypes^11^.

Ancient DNA, combined with results from GWAS on diet-related traits, provides a unique opportunity to study how multidimensional dietary phenotypes evolve through time because it allows direct observation of allele-frequency changes across successive populations experiencing distinct ecological and subsistence regimes, providing stronger inferences than those obtainable from modern-genome selection scans, where other evolutionary factors – such as admixture or drift–, may obscure or distort signals of past selection^13,14^. Previous work has begun to identify polygenic temporal changes in ancient DNA, but most studies have focused on individual loci, single traits, or broad GWAS panels without explicitly modelling genetic covariance among related endophenotypes^2,3,15–17^.

Holocene Europe provides a particularly suitable setting for studying dietary evolution, given its extensive ancient genomic record, the availability of GWAS data from European populations, and its well-characterized diversity of populations, environments, and subsistence strategies^18–21^. Over the last 10,000 years, Anatolian farming populations expanded across central and western Europe, while hunter-gatherer economies persisted longer in northern and eastern regions, maintaining a greater reliance on foraging, fishing, and aquatic resources. Later, the expansion of Steppe-associated groups contributed to the spread of mixed agro-pastoral economies. These asynchronous transitions reshaped food availability, resource exploitation, and subsistence practices, generating a mosaic of dietary environments across the continent.

In this study, we reconstruct the evolution of diet in European populations during the Holocene by integrating GWAS-derived dietary phenotypes with ancient genomic data. Rather than analysing individual diet-related traits in isolation, we infer latent dietary dimensions from patterns of genetic covariance and investigate their evolutionary dynamics through time and regions.

## Results

### Shared genetic architecture revealed four main clusters of diet-related traits

We initially compiled 94 GWAS datasets covering a broad spectrum of diet-related phenotypes, including anthropometric measures, body composition, metabolic, physical activity, and dietary behavioural traits (Supplementary Note 1). To characterize their shared genetic architecture, define biologically related families, and facilitate the identification of trait-specific genetic components for downstream analyses, we first evaluated GWAS quality using LDSC-derived diagnostics (Methods; Supplementary Note 3). LDSC intercepts were generally centred near 1, and most attenuation ratios were below 0.15 (Supplementary Figs. 6 and 7), supporting generally well-calibrated GWAS association tests, with the overall excess of association signal mainly attributable to polygenic effects rather than confounding^22–24^. Based on a combined assessment of LDSC diagnostics, SNP-heritability estimates (Supplementary Fig. 8), failed genetic-correlation analyses and dataset redundancy, 58 traits were retained for genetic-correlation and clustering analyses, with excluded traits detailed in Supplementary Table 2.

Pairwise bivariate LDSC analyses revealed widespread shared genetic architecture among the 58 retained traits. Of the 1,653 unique trait pairs, 1,295 (78.34%) showed nominally significant genetic correlations (P < 0.05), and 933 (56.44%) remained significant after Bonferroni correction (P < 3.0248 × 10⁻⁵; Supplementary Figs. 16 and 17). The 358 nominally non-significant pairs generally had estimates close to zero (median absolute *r*_g_ = 0.0274; 97.77% with |*r*_g_| < 0.10), although non-significance alone does not establish the absence of genetic correlation. To summarize this high-dimensional structure, genetic-correlation-derived dissimilarities were represented in a four-dimensional MDS configuration. The first two dimensions were selected for their interpretable trait groupings and preservation of local genetic neighbourhoods, hence used as input for a medoid-based genetic algorithm (GA). This optimization procedure searched candidate sets of representative traits, assigning each remaining trait to its nearest representative and favouring solutions with greater separation between representatives relative to within-cluster dispersion (Methods; Supplementary Note 4 and Supplementary Fig. 18). Among the tested cluster counts (k = 2–10), the highest observed Dunn-based fitness was obtained at k = 4 (Supplementary Fig. 19). Docosahexaenoic fatty acid (DHA) was the only trait subsequently reassigned from C4 to C3, being the only trait with an extreme intercluster genetic association whose ten strongest absolute genetic correlations provided greater support for the neighbouring cluster than its original cluster (Fig. 1; Supplementary Note 4). The resulting groups delineated a discrete partition within a continuous pattern of shared genetic architecture.

**Figure 1.**
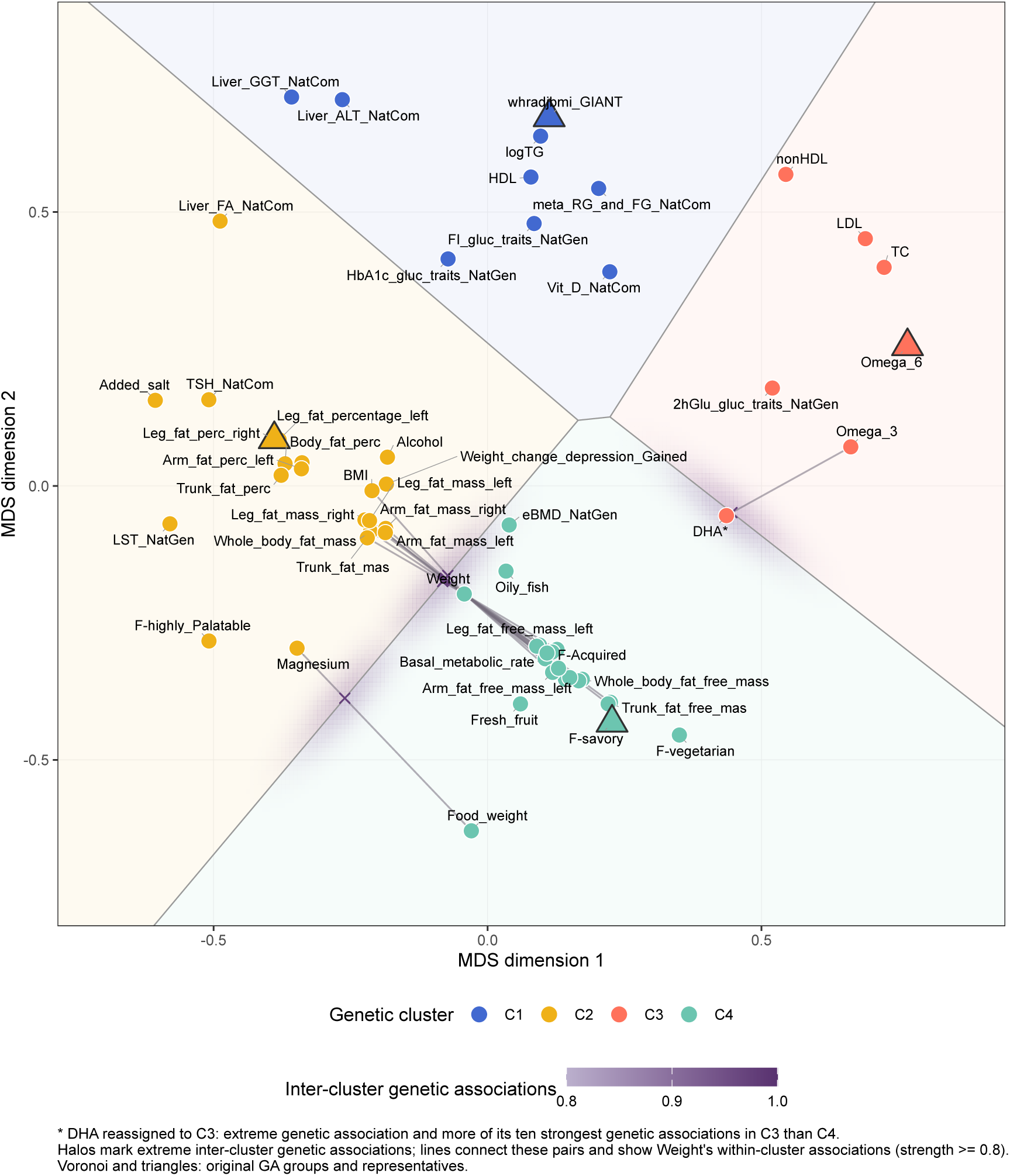
Shared genetic architecture defines four clusters of diet-related traits. MDS projection of shared genetic architecture among 58 diet-related traits, irrespective of effect direction. Points represent traits and are coloured by the cluster used in downstream analyses; triangles mark GA-selected medoids. Voronoi boundaries lie equidistant between neighbouring medoids. The clusters represent discrete groups within a continuous pattern of genetic relationships, with some traits retaining strong associations across boundaries. Halos highlight extreme intercluster genetic associations (absolute correlation ≥ 0.8), and lines connect the corresponding traits; equally strong intracluster associations involving Weight are also shown. Among traits with extreme intercluster associations, DHA* was the only case with greater genetic support in the neighbouring cluster and was therefore reassigned from C4 to C3. The remaining traits retained comparable or greater support within their original clusters. Boundaries show the original GA partition.

Diet-related clusters are represented in Fig. 1. The first cluster was composed mainly of metabolic-related traits, including glycaemic features, liver enzymes, HDL cholesterol levels and vitamin D, but also the waist-to-hip ratio (WHR). The second cluster was the most heterogeneous, grouping traits from multiple categories, including fat-related body composition and BMI, thyroid-stimulating hormone (TSH), dietary habits and preferences related to salt intake and highly palatable foods, leisure screen time (LST) that is a modern proxy of physical activity, and a psychiatric trait associated with weight change during depression. In contrast, the third cluster was more homogeneous and was primarily related to fatty acid metabolism, non-HDL cholesterol, and total cholesterol. Finally, the fourth cluster grouped traits related to fat-free mass body composition, estimated bone mineral density (eBMD), basal metabolic rate, and dietary habits and preferences associated with vegetarian- like foods, acquired food liking, savoury foods, and fruit and oily fish consumption.

To minimize overlapping genetic associations among related traits, our SNP ascertainment strategy prioritized trait-specific genetic components within each cluster, followed by LD clumping (Methods). This approach aimed to reduce the contribution of shared pleiotropic signals to downstream temporal and geographic analyses, allowing evolutionary trajectories to be characterized using markers more specific to each trait. Following this procedure, three traits retained fewer than 20 markers and were excluded from downstream analyses: Magnesium (13 markers) and weight gain during depression (10 markers) in C2, and Food weight (18 markers) in C4.

### Diet-related traits show distinct temporal trajectories in ancient European populations

Temporal trajectory analyses were performed on 938 unrelated ancient European individuals from Allentoft et al., spanning 400 to 10,000 years BP and excluding present-day samples. The final analytical dataset was defined according to the temporal weighting criteria described in Methods. It covers multiple European regions and time periods, as illustrated in Supplementary Fig. 3A. Consistent with this broad temporal and geographic transect, the dataset also captures substantial ancestry heterogeneity across ancient European populations (Supplementary Fig. 3B).

Using weighted Kendall’s *τ* , we quantified temporal changes in the proportion of risk alleles across the selected trait-specific markers, while accounting for uneven sampling across 500- year bins. Comparison with null distributions generated by randomization of risk alleles showed that 14 out of the 58 observed trajectories deviated from expectations under the null model (Supplementary table 3). These signals corresponded to approximately 44%, 32%, 14%, and 15% of the traits evaluated in clusters 1 to 4, respectively. After Benjamini–Hochberg (BH) correction within clusters, six diet-related traits retained significance at the false discovery rate (FDR) threshold (Fig. 2). Although the first cluster contained the highest proportion of significant traits, suggesting a possible concentration of signals in this biological group, permutation of P-values between clusters for multiple test correction indicated that this enrichment did not differ significantly from random expectation (one-tailed P = 0.12; Supplementary Fig. 21).

**Figure 2.**
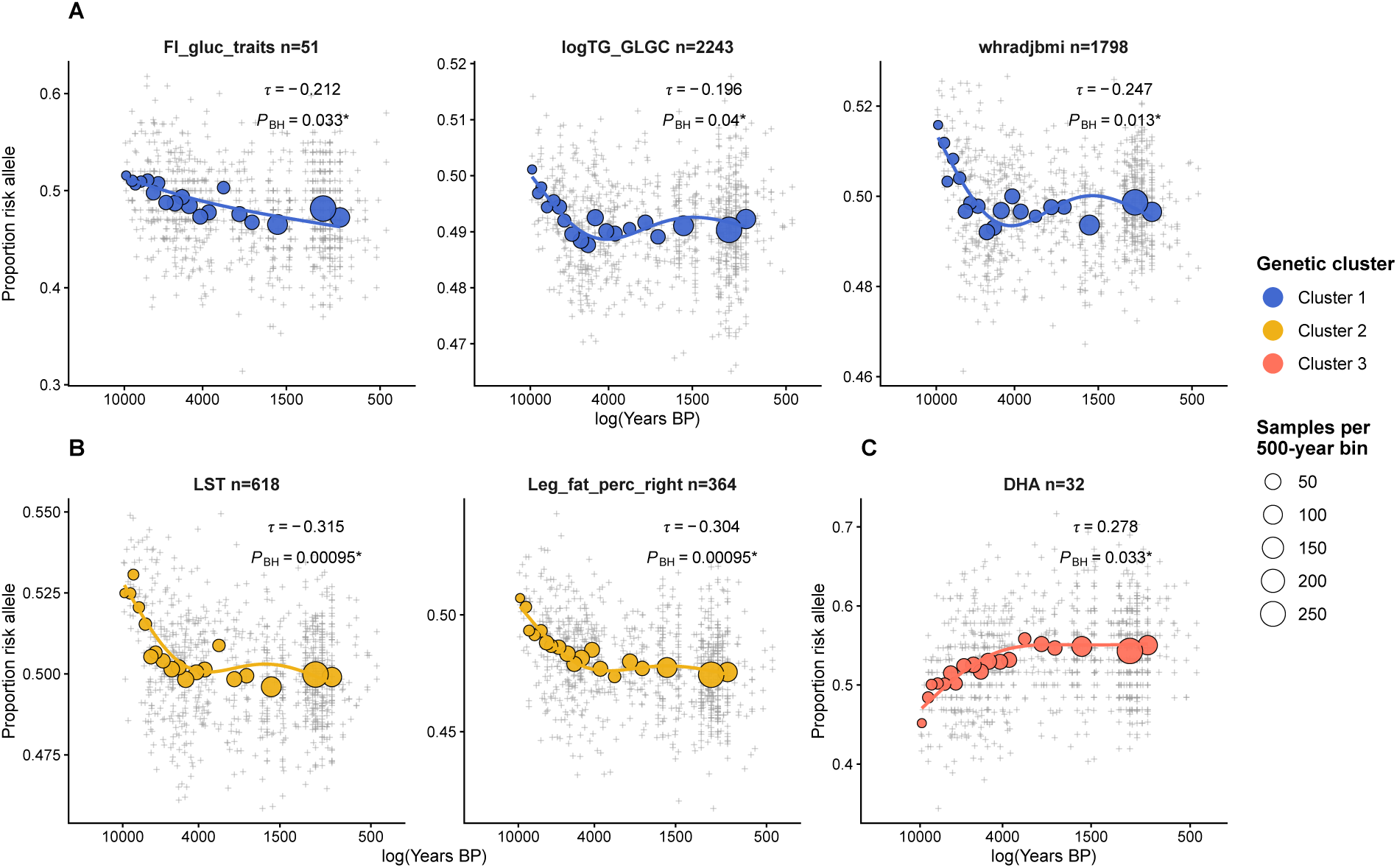
Pan-European temporal trajectories of BH-significant diet-related traits. Temporal trends in the proportion of risk alleles for the six diet-related traits that remained significant after Benjamini–Hochberg correction in the pan-European analysis. Trajectories were calculated using trait-specific genetic markers selected to reduce overlapping genetic associations among traits within each cluster, thereby limiting the contribution of shared pleiotropic signals (Methods). Panels show significant traits from cluster 1 (A), cluster 2 (B), and cluster 3 (C). The x-axis represents log-transformed years BP, spanning ancient samples from 400 to 10,000 years BP, with older periods shown on the left and more recent periods on the right. The y-axis represents the proportion of risk alleles. For each trait, the weighted Kendall’s *τ* and BH-adjusted P-value are shown; given the orientation of the x-axis, positive *τ* values indicate increasing risk-allele load toward the present, whereas negative *τ* values indicate decreasing risk-allele load toward the present. Traits are colored according to their diet-related domain. Smooth curves were fitted using a generalized additive model (GAM) with the same temporal-bin weights used in the weighted Kendall’s *τ* analysis, and are shown only as a visual representation of the trajectories. Circles represent temporal bins, with circle size proportional to the number of samples included in each bin. Legends indicate trait-domain colors and bin sample sizes.

The first cluster showed several significant temporal trajectories. Markers associated with WHR decreased rapidly from the Early Mesolithic (∼10,000 years BP) and then stabilized around 6,000 years BP (*τ* = −0.247, *P_BH_* = 0.013). A similar pattern was observed for triglyceride levels (logTG), although across a somewhat broader temporal interval (*τ* =−0.196, *P_BH_* = 0.04). In contrast, fasting insulin (FI) followed a more gradual decline extending from the Early Mesolithic to 400 years BP, without reaching a clear plateau by the Early Modern period (*τ* = −0.212, *P_BH_* = 0.033). Although HDL cholesterol did not survive FDR correction, its trajectory suggested an increase in associated markers from the Early Mesolithic to approximately 4,000 years BP, broadly corresponding to the Chalcolithic–Early Bronze Age transition in parts of Europe (*τ* = 0.18, *P* = 0.026).

In the second cluster, markers associated with right leg fat percentage showed a marked decrease during the Early Mesolithic, followed by gradual stabilization toward approximately 4,000 years BP (*τ* = −0.304, *P_BH_* = 9.5 × 10^−4^). A similarly strong pattern was observed for LST, which also decreased rapidly across the same broad interval (*τ* = −0.315, *P_BH_* = 9.5 × 10^−4^). LST showed moderate to high positive genetic correlations with fat-percentage traits. Additional body-composition traits within this cluster displayed concordant but weaker temporal patterns that did not survive FDR correction, including left leg fat mass (*τ* = −0.192, *P* = 0.0214) and whole-body fat mass (*τ* = −0.157, *P* = 0.0469). Together, these results suggest an overall decrease in markers related to adiposity until a plateau was reached around 4,000 years BP, near the Chalcolithic–Early Bronze Age transition. Further nominal signals were detected for alkaline phosphatase (*τ* = −0.162, *P* = 0.0394) and salt consumption habit (*τ* = 0.18, *P* = 0.0216).

In the third cluster, markers associated with docosahexaenoic fatty acid (DHA) increased from the Early Mesolithic until approximately 3,000 years BP, spanning the period after the Neolithic transition and into the Late Bronze Age (*τ* = 0.278, *P_BH_* = 0.033). DHA was the only trait in this cluster to show evidence of a non-random temporal trajectory.

No trait in the fourth cluster remained significant after FDR correction. Nevertheless, nominally significant decreases were observed for weight (*τ* = −0.203, *P* = 0.012), left leg predicted mass (*τ* = −0.216, *P* = 0.011), and right leg fat-free mass (*τ* = −0.143, *P* = 0.047). Here, predicted mass refers to impedance-derived estimates of body composition compartments, including water, lean mass, and fat. Overall, these trajectories also tended to stabilize around 4,000 years BP.

While our temporal analysis was restricted to ancient European populations from the last 10,000 years, we further investigated whether these traits exhibited differences in risk-allele representation between more distantly related human lineages, potentially reflecting divergent selective pressures over deeper evolutionary timescales. To this end, we compared the enrichment of risk-associated alleles for dietary traits between *Homo sapiens* and Neanderthals, using the Vindija Neanderthal genome and modern European variation data. Most traits showed no significant differences after multiple-testing correction. However, WHR showed significant enrichment (OR = 1.41, *P_BH_* = 0.028), indicating a higher proportion of WHR-increasing alleles in the Vindija Neanderthal genome than in modern Europeans.

### European demographic history does not fully explain the observed non-random signals

Because the dataset spans major episodes of migration and admixture across Europe^18,21^, we next evaluated whether the observed non-random trajectories could be explained by genome-wide demographic processes. To do so, we contrasted them against a conservative null model based on matched proxy alleles sampled from variants linked to protein-coding genes, which are expected to be especially sensitive to allele-frequency shifts driven by background selection and population history.

All traits that previously passed FDR correction remained statistically significant in this analysis, supporting the idea that the observed trajectories cannot be explained solely by demographic history. These included WHR (*P* = 0.001), logTG (*P* = 0.012) and FI (*P* = 0.011) from the first cluster; right leg fat percentage (*P* = 9.99 × 10^−4^) and LST (*P* = 9.99 × 10^−4^) from the second cluster; and DHA (*P* = 0.002) from the third cluster (Supplementary Fig. 28).

### Geographic partitioning reveals region-specific evolutionary trajectories for diet-related traits

Previously, temporal trajectories were evaluated across Europe as a whole. Because the individuals analyzed here span heterogeneous geographic origins, cultural contexts, and environmental conditions, regional differences may have contributed to distinct temporal patterns in diet-related traits. To test whether the evolutionary histories of diet-related traits differed across geographical regions, we implemented a genetic algorithm to identify geographic partitions that maximazed differences in temporal trajectories among aDNA samples (Methods; Supplementary Note 7 and Supplementary Fig. 29).

All traits that were statistically significant in the pan-European analysis showed geographic differentiation in their temporal trajectories (Fig. 4). For most traits, the inferred partition followed a broadly similar diagonal division between north-eastern and south-western Europe, approximately aligned with a Baltic–Black Sea axis. Across these traits, decreases in marker load were generally stronger in the eastern partition than in the western one. For FI and logTG in particular, western populations showed no temporal change since the Early Mesolithic, suggesting earlier stabilization in this region. In contrast, DHA from the third cluster showed a distinct geographic barrier orientation, extending diagonally from southern Spain to Finland and defining a less pronounced west–east partition of Europe. Under this configuration, DHA-associated alleles showed a sharper temporal increase in the western region. Overall, temporal changes were observed in both regions, although with different magnitudes and spatial configurations depending on the trait.

**Figure 3.**
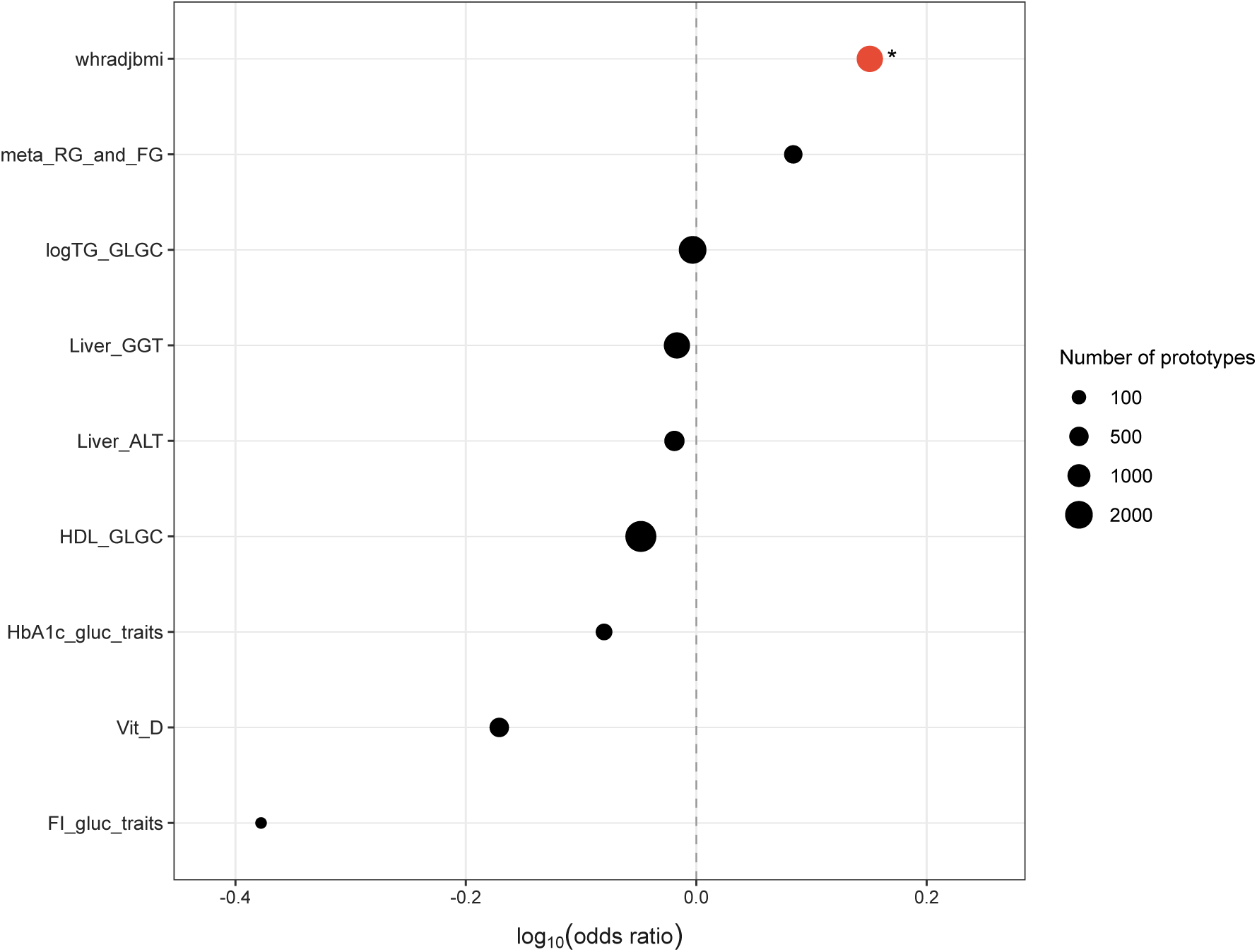
WHR shows significant enrichment of risk alleles in the Vindija Neanderthal lineage relative to modern Europeans. Enrichment analysis of risk alleles in the Vindija Neanderthal genome relative to modern European genetic variation for dietary traits in cluster 1. The x-axis represents the log- transformed odds ratio (log[OR]), where positive values indicate enrichment of trait-increasing risk alleles in the Vindija Neanderthal lineage relative to humans. Each point represents one tested trait, and point size is proportional to the number of independent markers included in Fisher’s exact test. Red indicates traits that remained significant after Benjamini–Hochberg correction within the cluster. Asterisks indicate BH-adjusted significance levels: *(*P_BH_* = < 0.05), **(*P_BH_* = < 0.01), and ***(*P_BH_* = < 0.001).

**Figure 4.**
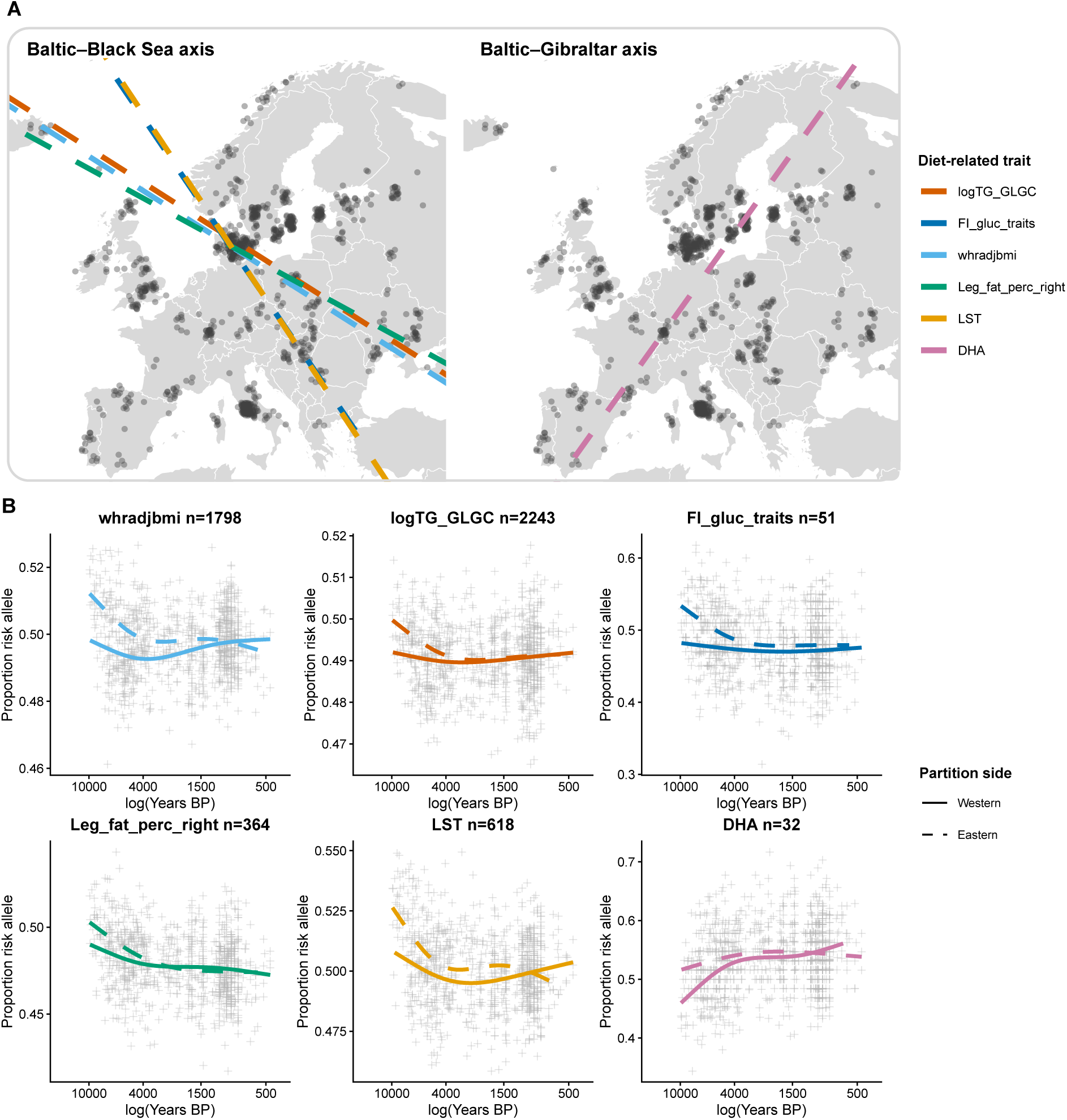
Geographic barriers reveal region-specific temporal trajectories of diet-related traits. Geographic partitioning and stratified temporal trajectories for the six diet-related traits that remained significant after Benjamini–Hochberg correction in the pan-European analysis. **(A)** European maps showing the aDNA samples included in the geographic analysis, represented as darker grey dots, and the geographic barriers inferred by the GA, represented as dashed lines. Two main barrier orientations were identified: a Baltic–Black Sea axis shared by five of the six significant traits, and a Baltic–Gibraltar axis identified for DHA. Dashed lines are colored according to the corresponding trait. Slight differences in line position reflect the approximate nature of the GA solutions for each trait. **(B)** Stratified temporal trajectories of the proportion of risk alleles for each trait according to the two sides of the inferred geographic partition. The x-axis represents log-transformed years BP, with ancient periods shown on the left and more recent periods on the right, and the y-axis represents the proportion of risk alleles. Smooth regression curves are shown separately for the western and eastern sides of each partition, using the same trait-specific colors as in panel A. Solid lines represent the western partition, whereas dashed lines represent the eastern partition. Unlike the pan- European trends, these smooths were not weighted and temporal bins were not displayed.

Inspection of the genetic fine-cluster composition of Mesolithic samples within each partition showed that the ancestry composition of the groups varied according to the inferred geographic barrier (Supplementary Figs. 30 and 31). For diet-related phenotypes WHR, FI, logTG, right leg fat percentage and LST, the geographic barrier followed the Baltic–Black Sea partition. In these phenotypes, the strongest eastern trajectories were observed among populations spanning approximately 10,000–4,000 years BP. Notably, the north-eastern Mesolithic groups often carried mixed eastern and western hunter-gatherer ancestry, whereas western Mesolithic groups were predominantly WHG-related. Consistently, western trajectories showed lower average marker load and weaker or absent temporal change. DHA displayed a different ancestral configuration: under its west–east partition, hunter-gatherer- associated groups from both western and eastern Europe began from lower allele frequencies, with the strongest increase subsequently observed on the western side of its partition. Overall, these patterns indicate that the regional differentiation identified by the genetic algorithm was already detectable before the later farmer- and steppe-related ancestry components observed in the final portions of several trajectories.

## Discussion

### Diet adaptation can be studied as interacting endophenotype systems

Diet has long been recognized as a major driver of human adaptation, yet it is rarely examined as a multidimensional biological system^1,3,5,13^. Most previous studies have focused on individual traits or loci, implicitly assuming that dietary adaptation can be deconstructed into a collection of largely independent evolutionary responses. By contrast, our framework groups genetically correlated phenotypes into endophenotype systems, allowing evolutionary change to be investigated in the context of shared genetic architectures. This approach is motivated by the observation that diet influences multiple interconnected physiological processes simultaneously, including energy storage, metabolic regulation, nutrient utilization and body composition. Such contextualization improves the interpretability of temporal trajectories in the presence of extensive pleiotropy, while enabling biologically informed multiple-testing correction that preserves statistical sensitivity within families of hypothesis (Methods). Importantly, significant trajectories reflect changes in trait-specific genetic components within these clusters, as defined by present-day GWAS, rather than direct reconstructions of ancient phenotypic values. Under this perspective, temporal changes are interpreted not as evidence for the evolution of isolated traits, but as signatures of modification in the broader biological systems that connect them.

### A coordinated adiposity-metabolism axis evolved through the Holocene

Our analyses identified coordinated temporal changes within Europe across multiple genetically correlated architectures related to adiposity, insulin regulation and lipid homeostasis, suggesting that adaptation may have acted on coordinated physiological systems rather than primarily on individual phenotypes. WHR-, triglyceride-, fasting insulin- and body-composition-related architectures broadly exhibited concordant temporal decrease in their trajectories, despite reflecting different physiological outcomes. Such concordance is biologically plausible because these phenotypes are linked by a common network regulating energy acquisition, storage and utilization. In particular, central adiposity is closely associated with insulin sensitivity, lipid partitioning and circulating triglyceride concentrations^25,26^, and changes in one component of this system often affect the others. Notably, fat-related body-composition traits predominated among the observed temporal trends, including non-FDR-significant declines in leg fat measures and whole-body fat mass.

The significant trajectory observed for leisure screen time (LST) further illustrates the advantages of an endophenotype-based perspective. Contemporary behavioural phenotypes are unlikely to represent direct targets of prehistoric selection, yet they can retain genetic correlations with metabolic architectures that were evolutionarily relevant in the past. LST has been genetically correlated with adiposity and metabolic traits^8^. In this sense, modern endophenotypes should not always be interpreted literally^27^. Rather, they can act as observable proxies for latent genetic architectures whose evolutionary origins predate the contemporary behaviours through which they are currently measured.

The coordinated nature of these trajectories is broadly compatible with previous ancient-DNA and polygenic-adaptation studies reporting evolutionary signals in anthropometric and metabolic traits, including declines in BMI-, WHR- and body-fat-associated genetic signals across West Eurasia^15,16^. Previous aDNA evidence of stabilizing selection on female WHR^17^, together with our finding of differential enrichment of WHR-increasing alleles in the archaic Homo-lineage comparison, further supports its evolutionary relevance across different timescales, consistent with previous evidence of significant heritability attributable to Neanderthal-informative variants for this phenotype^28^. However, these traits have generally been analysed and interpreted individually. By contrast, our results suggest that apparently distinct phenotypes can be viewed as partially overlapping windows into a shared adiposity- metabolism system.

One possible explanation for the predominance of significant signals within adiposity-related architectures lies in the biological role of adipose tissue itself. Adipose tissue is a highly plastic endocrine organ capable of responding to environmental conditions through changes in storage, mobilization and depot-specific regulation^29,30^, whereas lean tissues are often constrained by essential functions such as locomotion, thermogenesis and metabolic capacity^31,32^. Therefore, evolutionary responses affecting energy balance may be more readily accommodated through modifications in adipose regulation than through changes to more strongly constrained physiological systems. Such evolutionary responses may also have interacted with physiological trade-offs and cultural practices influencing energy availability and allocation, including food storage, sharing, mobility, seasonal dietary diversification and ritualized overfeeding during periods of abundance^18,33–36^.

Taken together, these observations suggest that Holocene subsistence transitions were accompanied by coordinated changes in energy storage and metabolic regulation. Importantly, such changes need not imply improved nutritional conditions. Early European farmers experienced nutritional, developmental and infectious challenges despite the adoption of agriculture, potentially contributing to their shorter osteological stature than expected from their genetic height scores^37^. Declining adiposity-related architectures are therefore not necessarily inconsistent with energetic stress. Instead, ecological and subsistence transitions may have altered the optimal balance between energy reserves and metabolic demands, favouring modifications in energy allocation, storage and utilization rather than simply promoting greater fat accumulation^13,16^.

### Multiple metabolic systems followed distinct evolutionary trajectories

Although the adiposity-metabolism system showed substantial temporal coherence, not all metabolic architectures followed the same evolutionary trajectory, supporting the perspective that different components of human metabolism appear to have responded to partially independent ecological, demographic and subsistence-related pressures.

In our analyses, the fatty-acid metabolism cluster exhibited a distinct pattern characterized by increasing DHA-associated alleles from the Early Mesolithic into the Late Bronze Age. The DHA trajectory is particularly notable because it diverges from the broader pattern observed for adiposity and metabolic regulation. This distinction is consistent with the idea that fatty-acid metabolism represents a partially independent biological module, influenced by dietary composition rather than primarily by mechanisms governing energy storage and partitioning. Previous aDNA studies have identified temporally variable selection at the FADS gene cluster, involved in the endogenous synthesis of long-chain polyunsaturated fatty acids, potentially reflecting changes in their dietary availability^2,38,39^. Changes in the balance between aquatic, animal and plant-derived foods during and after the Neolithic transition may therefore have shaped fatty-acid metabolism differently from systems involved in adiposity and glucose homeostasis. This is particularly notable for DHA, the only fatty-acid trait showing significant temporal changes when restricted to trait-specific variants.

In contrast, the decline in FI-increasing alleles may reflect metabolic changes more closely linked to the adiposity-related system. Indeed, genetic evidence connects FI to pathways involving adiposity, fat distribution, hepatic function and lipid metabolism^26^, consistent with our clustering analyses. The observed decline may therefore reflect changes in metabolic regulation associated with shifting energetic conditions. Such changes may also be compatible with improved insulin sensitivity or greater metabolic flexibility, alongside previous evidence of selection near insulin-regulatory loci in Neolithic Europeans^16^.

More broadly, the contrasting trajectories observed across clusters suggest that metabolism is organized into multiple interacting modules with overlapping but non-identical evolutionary histories. Some systems appear to have responded primarily to changes in energetic management and nutrient allocation, whereas others responded to shifts in dietary composition itself.

### Evolutionary histories were geographically heterogeneous

The temporal trajectories identified here were not distributed uniformly across Europe. Instead, several architectures displayed stronger or more gradual changes in north-eastern regions than in western populations. These differences are compatible with archaeological and genomic evidence indicating substantial regional variation in the timing and nature of subsistence transitions. Agriculture, pastoralism, foraging and aquatic-resource use existed in different combinations across Europe for millennia^18,20,40^, creating heterogeneous ecological contexts within which selection could operate.

Such patterns suggest that the same metabolic architectures may have experienced different selective regimes depending on local environmental conditions, subsistence practices and demographic histories. Climatic fluctuations, including the 8.2 ka and 4.2 ka events, provide further examples of how environmental conditions shaped subsistence strategies, with documented changes in animal management, diet and aquatic-resource exploitation^36,41^. In regions where agriculture was delayed or seasonally constrained, prolonged reliance on foraging, fishing and mixed economies may have maintained different balances between energy storage, nutrient availability and metabolic demand. The subsequent convergence of regional trajectories during later prehistory is also consistent with increasing connectivity, admixture and the spread of Steppe-associated populations.

Notably, DHA showed a distinct geographic partition, with temporal increases in both regions, but more pronounced on the western side of this partition, in contrast to the previous patterns. Previous evidence indicates that PUFA-associated alleles at FADS1/2 were already relatively common in early Middle Eastern farmers and Caucasus hunter-gatherer-related populations

before the Neolithic expansion across Europe, subsequently increasing across European hunter-gatherer and Anatolian farmer ancestral backgrounds^2^. Such ancestry-related differences in baseline allele frequencies, potentially combined with changes in dietary composition, may have contributed to the observed geographic contrast.

Consequently, our results suggest that dietary adaptation was not a uniform pan-European process but rather as a mosaic of regionally distinct evolutionary responses.

## Conclusions

Our findings provide evidence that dietary adaptation may be better understood as the coordinated evolution of several interconnected physiological systems. These changes were geographically heterogeneous, with stronger and more gradual shifts in north-eastern Europe, consistent with regionally distinct ecological, demographic and subsistence histories. Throughout the Holocene, major migrations and admixture events reshaped the genetic backgrounds on which selection operated^19,21^, while changes in diet, settlement patterns, pathogen exposure and population density created novel selective environments. Together, these interacting processes may have contributed to the diverse evolutionary trajectories of genetic architectures underlying metabolic regulation and energy balance across Europe.

## Methods

### Data collection

#### Diet-related traits

Genome-wide association study (GWAS) summary statistics and meta-analyses from traits associated to the diet phenotype were obtained from publicly available databases and consortia, including the GWAS Catalog, UK Biobank, the Genetic Investigation of ANthropometric Traits (GIANT), the Meta-Analyses of Glucose and Insulin-related traits Consortium (MAGIC), the Global Lipids Genetics Consortium (GLGC), the Program in Complex Trait Genomics, and the Genetic Factors for Osteoporosis (GEFOS) consortium. In total, 94 diet-related complex traits were initially compiled for the present work (Supplementary table 1). These traits spanned six main domains: metabolism, anthropometric measures, body composition, physical activity, psychiatric traits, and dietary composition and preferences (Supplementary Fig. 1). Because GWAS statistical power depends strongly on cohort size, the final collection included studies with substantial variability in sample size, with the largest cohorts generally corresponding to more recent meta-analyses (Supplementary Fig. 2).

#### Ancient genomes

Genotype data for 962 ancient individuals were analysed from Allentoft et al. (2024), who generated and curated a post-glacial western Eurasian ancient DNA dataset imputed together with previously published individuals^18^. The initial ancient European subset considered here spanned approximately 200 to 15,000 years before present (BP). This subset comprised heterogeneous ancient groups spanning western and eastern hunter-gatherers, early and later Neolithic farmer-associated populations, post-Neolithic groups from western, northern and eastern Europe, post-Neolithic eastern Mediterranean/Levant-associated groups, and early and later steppe-related pastoralist groups, with Faroese individuals retained as an additional regional group. These group labels were used for the MDS visualization shown in Supplementary Fig. 3 and provide a broad summary of the demographic layers represented in the dataset. The dataset had already been imputed with GLIMPSE v.1.0.0 and includes variants from both the 1000 Genomes Project and the 1240K array. In addition, the archaic genome of the Vindija Neanderthal was obtained from the Max Planck Institute^42^.

### Modern European populations

European individuals from the 1000 Genomes Project Phase 3 release^43^ were used as the modern reference panel. We included the five European populations CEU, TSI, FIN, GBR and IBS, comprising 503 individuals in total, and retrieved genotype data from the Phase 3 integrated call set available through the 1000 Genomes FTP repository (https://ftp.1000genomes.ebi.ac.uk/vol1/ftp/release/20130502/). The full Phase 3 integrated call set contains more than 79 million variant sites, including SNPs, indels and other variant classes.

### Quality control

#### GWAS data

Linkage Disequilibrium Score Regression (LDSC)^22,44^ was applied to the selected GWAS in order to estimate the proportion of phenotypic variance explained by their genetic architectures. Precomputed LD scores for European populations based on the 1000 Genomes Project Phase 3 reference panel were used throughout the analysis, covering up to 1,217,311 genome-wide variants. To harmonize variants across GWAS and LDSC reference files, genomic coordinates were first lifted over to hg19 when necessary. Variants were then

matched by physical position using the join -1 1 -2 1 command. A chr:pos identifier was used to minimize discrepancies in SNP identifiers across studies and to maximize recovery of overlapping variants. Finally, pairwise LDSC bivariate analyses were performed using ldsc.py with the arguments --rg trait1,trait2, --w-ld-chr eur_w_ld_chr/, and --ref-ld-chr eur_w_ld_chr/.

Both univariate and bivariate cross-trait LDSC regressions were performed. After harmonization with the European LDSC reference panel, most traits retained approximately 1 million variants, and all GWAS retained more than 300,000 variants. This exceeded the 200,000-variant warning threshold described in the LDSC documentation, below which regression results may indicate problematic merging with LD scores or yield unreliable estimates. The distribution of retained variants across GWAS is shown in Supplementary Fig. 5.

GWAS suitability for genetic-correlation and clustering analyses was assessed using LDSC intercepts, attenuation ratios and SNP-heritability estimates. Intercepts and attenuation ratios were inspected to assess calibration and distinguish inflation attributable to polygenic signal from other sources^22^. Univariate intercepts were evaluated relative to their expected value of approximately 1^22^, and attenuation ratios >0.20 were used to flag potentially problematic GWAS, informed by previous applications of LDSC diagnostics^23,24^. GWAS selection combined these diagnostics with the identification of failed genetic-correlation analyses, a study-specific SNP-heritability threshold of 0.05, and the removal of redundant datasets, prioritizing those with more favourable LDSC metrics and greater statistical power. The rationale and supporting diagnostics are detailed in Supplementary Note 3 and Supplementary Table 2.

### Ancient genomes

Ancient individuals were further processed according to the quality-control recommendations provided by Allentoft et al. Variants were excluded when the imputation INFO score was < 0.8 or the minor allele frequency was < 0.01, and individual genotypes with genotype probabilities < 0.8 were treated as missing data. Only unrelated individuals without sample-level quality-control flags were retained. Variant filtering was performed using bcftools v1.19 with bcftools view -S ${sample_file} -i ’MAF >= 0.01 & INFO >= 0.8’ , and low-confidence genotypes were set to missing using bcftools +setGT -Oz -- -t q -n . -e ’GP[*] >= 0.8’.

To reduce bias in temporal trajectory analyses arising from uneven sampling across time^45^, individuals were grouped into 500-year temporal bins, and each individual was assigned a temporal weight of *w_i_* = 1/*n_b_*, where *n_b_* represents the number of individuals within its bin. This weighting scheme prevented densely sampled periods from disproportionately influencing temporal trends. The initial ancient subset spanned approximately 200–15,000 BP.

However, individuals older than 10,000 BP were sparsely represented, with some 500-year bins containing only a single individual, resulting in disproportionately large individual weights (Supplementary Fig. 4). To avoid temporal trends being driven by these poorly represented intervals, samples older than 10,000 BP were excluded. Similarly, individuals more recent than 400 BP and present-day samples were excluded to avoid anchoring temporal patterns to recent genetic structure. Downstream analyses were therefore restricted to unrelated ancient individuals dated between 400 and 10,000 BP, comprising a final dataset of 938 ancient genomes.

### Diet-related traits clustering

Several clustering strategies were initially explored, ranging from traditional dimensionality reduction approaches to more computationally intensive methods. When conventional approaches such as PCA and k-means were applied directly to GWAS summary statistics, traits clustered primarily by consortium or study source rather than by biological meaning (see Supplementary Note 4), likely reflecting shared cohorts, phenotype definitions, and residual study-specific structure rather than genuine biological relationships. Traits derived from the UK Biobank accounted for most of the observed variance and dominated the clustering structure, while biological heterogeneity across traits collected from different sources was not captured, leading to their aggregation into a single group. As genetic correlations provide a more robust representation of shared genetic architecture across traits, a pairwise matrix of genetic correlations was calculated using LDSC bivariate regression analysis for the final set of 58 diet-related traits.

Although genetic correlations provide an informative representation of shared genetic architecture, the resulting pairwise matrix does not by itself define an objective clustering solution. To enable quantitative comparison of alternative trait partitions, absolute genetic correlations were transformed into a dissimilarity matrix defined as 1 - |*r*_g_|. Using absolute values allowed both positive and negative genetic correlations to contribute to trait grouping, under the assumption that agonistic and antagonistic relationships may both reflect shared biological or evolutionary constraints. This dissimilarity matrix was projected using Multidimensional Scaling (MDS) with the R package smacof, and the first two dimensions of the embedded space were used as input for a genetic algorithm (GA) implementing a k-medoids-based clustering strategy to identify the optimal trait partition. Details on the MDS implementation and parameter selection are provided in Supplementary Note 4.

### Genetic algorithm-based k-medoids clustering

To identify groups of diet-related phenotypes with shared genetic architectures, we implemented a genetic algorithm, a population-based metaheuristic optimization approach that stochastically explores candidate clustering solutions and iteratively improves them through nature-inspired strategies (Supplementary Fig. 18). Therefore, it searched over sets of *k* representative traits in the D1–D2 projection of a four-dimensional MDS configuration. Representatives were selected from observed traits rather than calculated as arithmetic centroids, yielding a medoid-based partition. Each candidate solution comprised *k* distinct representatives, and the remaining traits were assigned to their nearest representative using Euclidean distances. Clustering quality was evaluated using a Dunn index that favoured separation between representatives relative to within-cluster dispersion. Candidate solutions evolved through elitist selection, crossover and mutation. We evaluated *k* = 2–10 and selected *k* = 4, which yielded the highest observed fitness among the tested values. The fitness definition, evolutionary operators, algorithm parameters and choice of MDS projection are detailed in Supplementary Note 4.

The GA-derived partition was subsequently reviewed for extreme intercluster genetic associations (absolute genetic correlation ≥ 0.8). For traits involved in these associations, reassignment required greater representation of the candidate neighbouring cluster than the original cluster among the trait’s ten strongest absolute genetic correlations. DHA was the only trait meeting this criterion and was reassigned from C4 to C3 for downstream analyses; the remaining traits retained comparable or greater support within their original clusters. This post-clustering refinement is detailed in Supplementary Note 4

### SNP ascertainment

After clustering diet-related traits, we designed a strategy to construct trait-specific SNP libraries while accounting for LD structure and shared within-cluster genetic architecture. Because pleiotropic markers may be subject to antagonistic pressures across traits, their inclusion could reflect divergent evolutionary histories. We therefore restricted SNP ascertainment to markers with an approximately independent evolutionary history within each biological group.

As a first step, we implemented a Python-based pipeline to harmonize GWAS summary statistics across studies. Quality control included exclusion of structural variants, multiallelic and non-biallelic markers, and variants located on non-autosomal chromosomes. SNPs were then matched across datasets using allele information and genomic coordinates to ensure consistent variant representation between studies. We subsequently generated a cluster-specific dataframe summarizing shared and trait-specific variants within each cluster. Associations were retained at a suggestive P-value threshold of P < 1 × 10⁻⁵ to capture candidate trait-associated variants for downstream aggregation, rather than to define individually genome-wide significant loci. This is consistent with previously proposed suggestive GWAS thresholds based on the effective number of independent tests^46^. Moreover, conventional Bonferroni-derived genome-wide significance thresholds can be conservative, and biologically constrained sub-significant GWAS signals may contain potentially true-positive loci for further investigation^47,48^.

Next, uniquely associated SNPs per trait were intersected with the ancient DNA dataset, defining the initial set of variants eligible for temporal trajectory analyses. These variants were subsequently intersected with present-day European data from the 1000 Genomes Project. LD was not inferred from ancient samples, as their heterogeneous temporal distribution could obscure true shared evolutionary history among markers and reduce statistical power. Instead, clumping was performed on the intersected GWAS–aDNA–1000G set using European 1000G genotypes with PLINK v1.9b5 (--geno 0.8, --clump-p1 1e-05, --clump-p2 1e-05, --clump-r2 0.2, --clump-kb 500). Traits retaining at least 20 independent markers after clumping were considered sufficiently polygenic to proceed to downstream analyses, consistent with a previously used threshold^16^. Traits excluded based on this criterion are listed in Supplementary table 2.

Importantly, hypervariable genomic regions such as the MHC and/or regions with complex LD patterns (structural variation, etc.) were not included in this clumping. Only the most significant SNP was considered in each of the following genomic segments (hg19): chr6:25,500,000– 33,500,000, chr8:8,135,000-12,000,000 and chr17:40,900,000-45,000,000.

### Temporal trend analysis

Because a linear relationship between trait values and time was not assumed, consistent with the expected evolutionary dynamics of complex traits shaped by heterogeneous stages of selection^11^, temporal trends were assessed using a weighted Kendall’s *τ* correlation. To account for uneven temporal sampling in ancient DNA datasets, which can otherwise allow densely sampled periods to disproportionately influence temporal trends^45^, we incorporated the temporal weights derived from 500-year bins, as described above. Specifically, we quantified the association between the proportion of risk alleles and sample age (years BP) as:

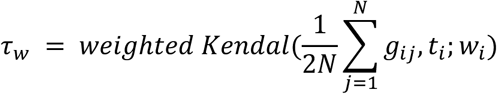

where *g_i_*_j_ ∈ {0,1,2} denotes the genotype of individual *i* at SNP *j*, *N* is the number of SNPs included for the trait, and *t_i_* represents the age of the sample in years BP, and *w_i_* = 1/*n_b_* is the temporal weight assigned to individual *i*, with *n_b_* denoting the number of individuals in its corresponding 500-year bin. Tied pairs were excluded from the denominator when computing weighted Kendall’s *τ*, making the resulting statistic equivalent to a weighted version of Goodman–Kruskal’s gamma. This approach allows assessment of both the direction and strength of the temporal association.

To evaluate whether the observed temporal trends exceeded expectations under randomized risk-allele assignments, we generated 10,000 null replicates by randomizing risk-allele assignments and recalculating weighted Kendall’s *τ* at each iteration. One-tailed Monte Carlo empirical P-values were derived from the resulting null distributions according to the direction of the observed effect, using the upper tail for positive *τ* values and the lower tail for negative *τ* values. These P-values were used to identify traits showing temporal signals that deviated from expectations under the assumption of no association between the reference allele and the phenotype. This approach is conservative because all genetic markers were treated equally and effect size estimates from present-day populations were not applied to ancient samples.

To account for the dependence among temporal tests arising from shared genetic architectures, traits were clustered using absolute genetic correlations, capturing both concordant and antagonistic pleiotropic relationships. As pleiotropy can induce correlations among trait-associated test statistics, genetically correlated traits were expected to exhibit more strongly dependent temporal trajectories within clusters than between clusters, implying redundancy among hypotheses and a smaller effective number of independent tests. More broadly, incorporating biologically informed group structures into multiple-testing frameworks has been shown to improve statistical power by exploiting differences in signal enrichment across groups^49^. We therefore treated endophenotype clusters as approximate families of related hypotheses and applied the Benjamini–Hochberg (BH) procedure separately within each cluster, which accommodates independence and certain forms of positive dependence^50,51^. As families were defined independently of temporal test significance and all clusters were tested, no additional family-selection adjustment was required. This approach targets the average false discovery rate across endophenotype families, rather than the pooled false discovery rate across all traits^52^.

### Assessment of demographic susceptibility using a genome-wide empirical background

Traits that remained statistically significant after FDR correction in the temporal analysis exhibited non-random temporal patterns. However, migration, admixture and population substructure can also generate systematic allele-frequency changes across ancient populations, potentially confounding the interpretation of evolutionary signals. To assess whether the observed trait-associated trajectories exceeded genome-wide variation potentially influenced by these demographic processes, we constructed an empirical null distribution from variants linked to protein-coding genes.

Because purifying selection in coding regions removes deleterious and slightly deleterious variants, linked markers may also be affected through background selection. As a result, SNPs linked to protein-coding genes may be particularly sensitive to genome-wide differences in allele frequencies among populations, reflecting the combined influence of demographic history and linked selection. For this reason, these variants were used to construct an empirical null set against which to evaluate the observed temporal trajectories. This set was recovered using variant coordinates from ancient samples and a gene-only BED file derived from the GENCODE hg19 release 47 lift37 annotation^53^. Only annotations with confidence levels of 2 and 3 were selected. The nearest protein-coding genes were identified using bedtools closest (v2.30.0), and variants located within genes or within 50 kb of their boundaries were retained.

For each tested trait, proxy marker sets were sampled from the genome-wide null set by matching allele frequency and the number of markers used to calculate the observed proportion of risk alleles. For each replicate, the proportion of proxy alleles was calculated across ancient individuals, and the temporal association between proxy allele proportion and sample age was estimated using the same weighted Kendall’s *τ* framework applied to the observed trait-associated markers. This procedure was repeated across 1,000 random samplings of the null SNP set, generating a trait-specific empirical null distribution of weighted Kendall’s *τ* values reflecting temporal variation in the genome-wide background. The observed weighted Kendall’s *τ* was then compared with this distribution, and one-tailed Monte Carlo empirical P-values were calculated from the upper or lower tail according to the sign of the observed effect.

### Geospatial trend analysis

A temporal trend observed at the pan-European level could partly arise from spatial heterogeneity. To identify geographic regions exhibiting highly differentiated temporal trajectories for a given trait, we developed a genetic algorithm that iteratively optimized candidate linear geographic barriers through stochastic selection, recombination and mutation

(Supplementary Fig. 29). Each candidate barrier partitioned ancient samples into two geographic groups. The fitness of each solution was defined by an F-like statistic, computed as the ratio of inter-group to intra-group variance in temporal trajectories. In this way, the algorithm favoured partitions that maximized differences between regions while minimizing dispersion within each region.

To avoid poorly comparable solutions driven by highly unequal sample sizes, an additional penalty term was applied to strongly imbalanced partitions. This constrained the search toward more balanced groupings while preserving the objective of maximizing regional differentiation. Candidate solutions evolved through tournament-based parent selection and evolutionary operators, as detailed alongside the fitness definition and algorithm parameters in Supplementary Note 7.

### Selective pressures in homo lineages vs modern humans

To assess whether selective pressures acting on diet-related traits may extend deeper in evolutionary time, we compared the enrichment of trait-associated derived alleles between the Vindija Neanderthal genome and *Homo sapiens*, using modern European variation data. Ancestral allele information was retrieved from the 1000 Genomes Project Phase 3, where ancestral states were inferred from alignment to *Pan troglodytes*. GWAS variants associated with each tested trait were then overlapped with the Vindija genome using physical genomic coordinates to obtain the set of available markers. For each trait, a contingency table was constructed based on the number of derived alleles present in the Vindija Neanderthal lineage and in modern humans that either increased or decreased the trait value. Fisher’s exact test was then applied to evaluate differences in the corresponding odds ratios (OR). Because the expected direction of enrichment depends on the temporal trajectory observed in the ancient DNA analysis, the one-tailed P-value from Fisher’s exact test was selected according to the directionality of the corresponding temporal effect. For each trait, the alternative hypothesis was chosen to test whether the enrichment of risk-increasing alleles in the Vindija Neanderthal lineage relative to humans was consistent with the direction of temporal change observed in the ancient DNA dataset. After selecting the direction-specific one-tailed P-value for each tested trait, resulting P-values were adjusted using the Benjamini–Hochberg procedure within each trait cluster.

## Acknowledgements

We thank all colleagues and researchers who provided valuable feedback and constructive discussions throughout the development of this study.

## Code availability

The custom code developed for data processing, statistical analyses and visualization in this study is publicly available on GitHub at https://github.com/ccsbio/dietary-endophenotype-evolution

