## Supplementary Material for "Beyond single-trait analysis: ancient genomes reveal coordinated evolution of dietary endophenotypes in Holocene Europe"

### Supplementary Materials

#### Supplementary Note 1. Data collection of diet-related traits

The initial GWAS collection was designed to capture multiple dimensions of the diet-related phenotype, including metabolic, anthropometric, body-composition, behavioural and dietary-composition traits. Publicly available GWAS summary statistics and meta-analyses were prioritized when they corresponded to large, well-powered studies, recent releases, or widely used consortia resources. In total, 94 traits were initially compiled before downstream quality control. The distribution of studies across the six diet-related domains is shown in Supplementary Fig. 1, and the sample-size distribution of UK Biobank and non-UK Biobank studies is shown in Supplementary Fig. 2. The complete list of GWAS initially considered is provided in Supplementary Table 1.

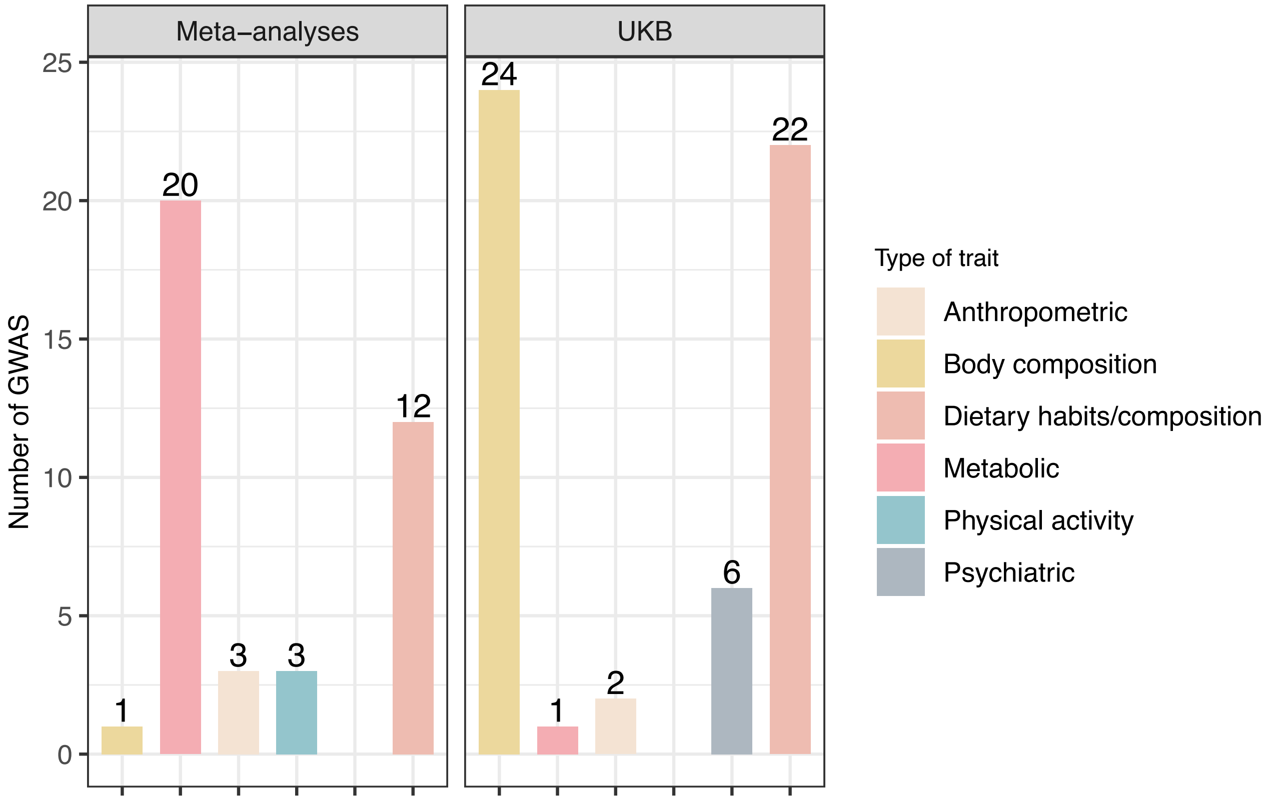


**Supplementary Fig. 1. Distribution of GWAS studies by diet-related domain.**
Number of GWAS traits initially compiled for each of the six diet-related domains considered in this study: metabolism, anthropometric measures, body composition, physical activity, psychiatric traits, and dietary composition and preferences. These domains were used to organize the initial GWAS collection before LDSC-based quality control.

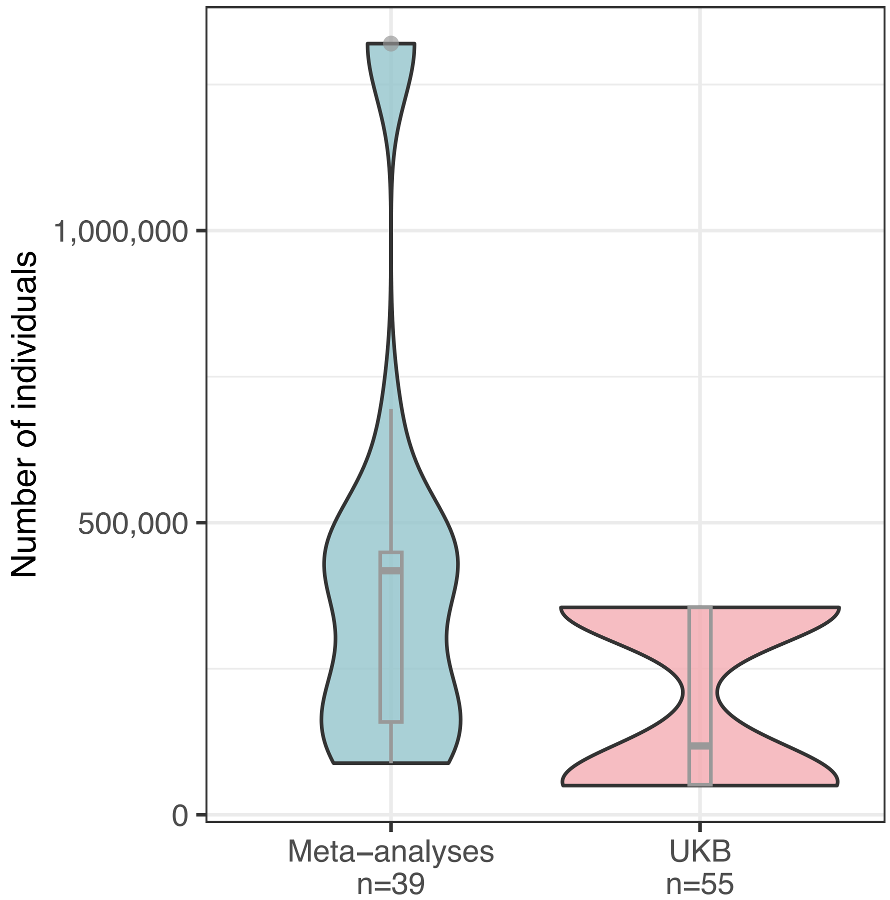


**Supplementary Fig. 2. Sample size distribution of UKB and non-UKB GWAS studies.** Distribution of GWAS sample sizes among the 94 initially compiled diet-related traits. Studies are classified according to whether the GWAS was based primarily on UK Biobank data or on non-UK Biobank cohorts/consortia. Sample size variability reflects differences in trait availability, study design and cohort scale across diet-related domains.

#### Supplementary Note 2. Ancient DNA dataset, quality control and temporal sampling

Additional visual summaries of the ancient DNA dataset are provided to complement the sample description and filtering criteria reported in the Methods. Supplementary Fig. 3 summarizes the geographic and temporal distributions, together with the genetic structure of the 962 ancient individuals initially considered: panel A shows the spatial distribution of samples across Europe, with points coloured by sample age, whereas panel B shows the MDS of genetic distances coloured by the original `Cluster.group` metadata label. Supplementary Fig. 4 illustrates the uneven temporal sampling of the dataset after excluding individuals more recent than 400 BP, but before applying the upper age cutoff of 10,000 BP. Panel A shows the number of individuals per 500-year bin, including poorly represented intervals extending back to 15,000 BP, whereas panel B shows the distribution of temporal weights, highlighting the disproportionately large weights assigned to sparsely represented periods. Together, these distributions support the exclusion of samples older than 10,000 BP from downstream temporal analyses.

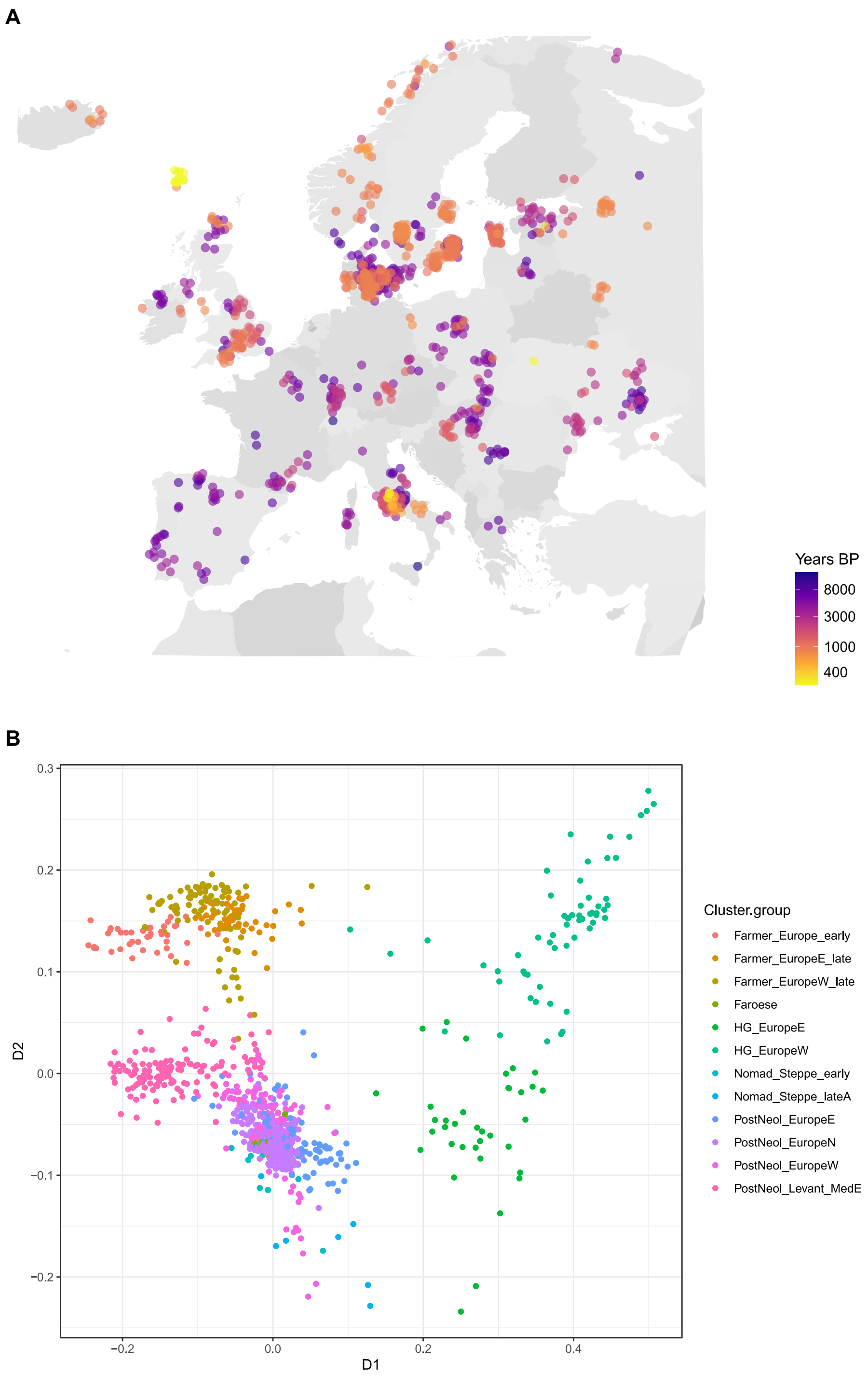


**Supplementary Fig. 3 | Geographic, temporal and genetic structure of the initial ancient DNA dataset.**

**A**) geographic distribution of the 962 ancient individuals initially considered from Allentoft et al. (2024), spanning approximately 200–15,000 years BP. Points represent ancient samples and are coloured according to sample age, showing the temporal and spatial coverage of the dataset across Europe.

**B**) MDS visualization of genetic distances among the same 962 ancient individuals. Points are coloured according to the original `Cluster.group` metadata label from Allentoft et al. The MDS summarizes broad population structure across the initial ancient dataset.

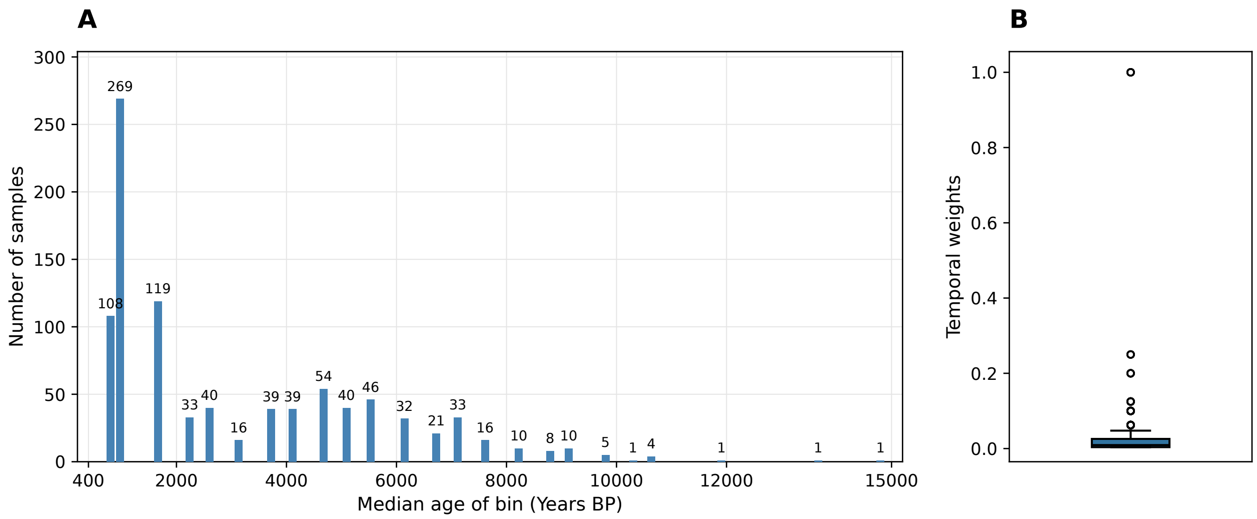


**Supplementary Fig. 4 | Temporal sampling and weight distribution of the ancient DNA dataset.**

**A)** Number of ancient individuals per 500-year bin after excluding samples more recent than 400 BP, but before applying the upper age cutoff of 10,000 BP. The low sample counts in the oldest intervals motivated their exclusion from downstream temporal analyses.

**B)** Distribution of temporal weights across the same dataset. Weights were defined as the inverse of the number of individuals per 500-year bin, resulting in disproportionately large weights in sparsely represented periods, with single-sample bins receiving a weight of 1.

#### Supplementary Note 3. GWAS harmonization and LDSC quality control

This section provides additional details on the harmonization and LDSC-based quality-control procedure applied to the 94 initially compiled GWAS summary statistics. Prior to LDSC regression, variants from each GWAS were matched to the European LDSC reference panel using physical genomic coordinates. A `chr:pos` identifier was generated to reduce inconsistencies due to rsID updates or missing SNP identifiers across studies, and overlapping variants were recovered using the Unix `join -1 1 -2 1` command.

Summary statistics were standardized using `munge_sumstats.py`. Parameters were adapted to the information available in each GWAS, using either `--N` or `--N-col` depending on whether sample size was reported globally or per variant. Effect-size columns were specified with `--signed-sumstats ${sumstat_col},${null_value}`, with the null value set according to the type of statistic provided by each study, including beta, log odds, odds ratio or Z-score. Additional parameters included `--chunksize 500000` and `--merge-alleles eur_w_ld_chr/w_hm3.snplist`, restricting analyses to HapMap3 variants represented in the LDSC reference panel.

After harmonization and munging, most analyses included approximately one million variants, and all GWAS retained more than 300,000 variants (Supplementary Fig. 5). Suitability for downstream analyses was assessed using LDSC intercepts, attenuation ratios and SNP-heritability estimates. The univariate intercept is expected to be approximately 1, although genomic-control correction can produce values below 1[1]. The attenuation ratio, defined as (intercept − 1)/(mean $\chi^2$ − 1), estimates the proportion of inflation attributed by the LDSC model to sources other than polygenic signal. A value of 0.20 was used as a pragmatic diagnostic reference, following previous applications assessing compatibility between GWAS summary statistics and LD reference panels[2,3].

For visualization, LDSC intercepts, attenuation ratios and SNP-heritability estimates were summarized per trait by averaging estimates across pairwise analyses, prior to quality-control filtering. Quality-control criteria were assessed using the underlying estimates.

GWAS selection was based on a combined assessment of LDSC diagnostics, SNP-heritability estimates, successful genetic-correlation analyses and redundancy among datasets. Two traits with failed genetic-correlation analyses were identified. A study-specific SNP-heritability threshold of 0.05 was also applied, with traits excluded if any estimate fell below this threshold in the pairwise analyses. This criterion was intended to reduce potential instability associated with low heritable signal, rather than represent a minimum requirement of LDSC. In a descriptive comparison after removing the two traits with failed analyses, pairs involving at least one SNP-heritability estimate below 0.05 had a higher median genetic-correlation standard error than the remaining pairs (0.074 versus 0.025). This comparison supported the motivation for the filter but did not establish 0.05 as an optimal threshold.

For redundant GWAS datasets, selection prioritized those with more favourable LDSC diagnostics and greater statistical power. Overall, 36 of the 94 initially compiled traits were excluded through the combined QC and redundancy assessment, leaving 58 traits for genetic-correlation and clustering analyses. None of the retained traits had a mean attenuation ratio of 0.20 or higher. Intercept and attenuation-ratio distributions are shown in Supplementary Figs. 6 and 7, respectively, and SNP-heritability distributions stratified by UK Biobank and non-UK Biobank studies are shown in Supplementary Fig. 8. A further three traits were excluded from downstream evolutionary analyses because they retained 20 or fewer independent markers after LD clumping. The complete list of excluded traits and their exclusion reasons is provided in Supplementary Table 2.

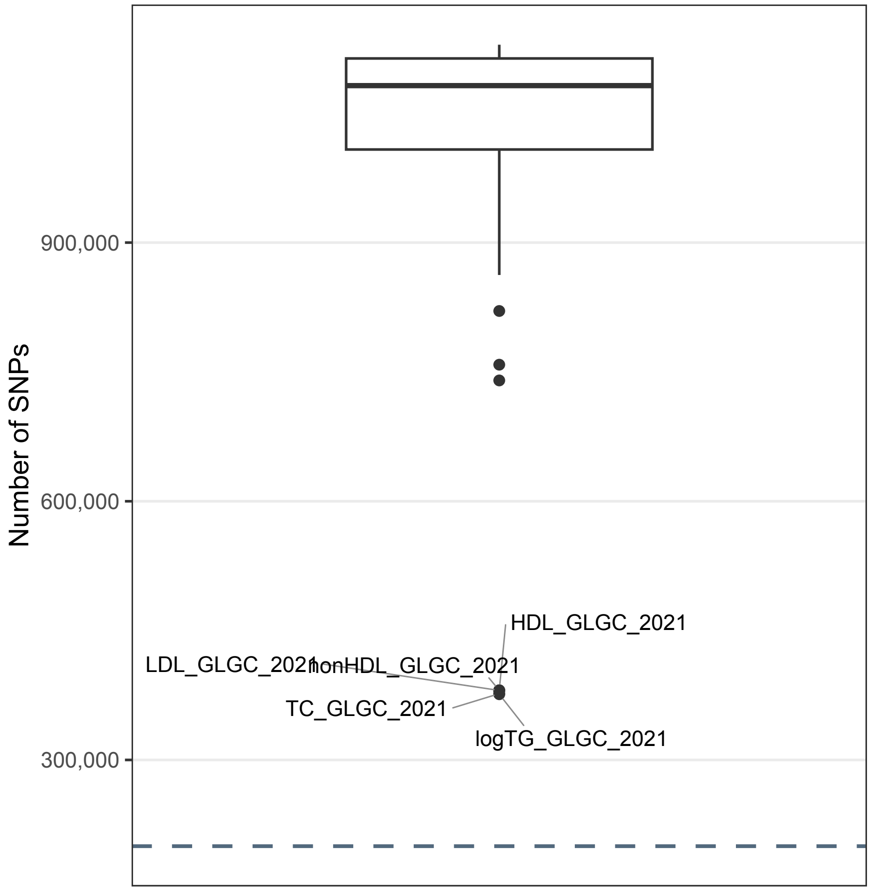


**Supplementary Fig. 5 | Number of genetic markers retained per GWAS for LDSC regression.**

Distribution of the number of variants retained for LDSC regression across the 94 initially compiled GWAS datasets after harmonization, munging and merging with the European HapMap3 SNP list used by LDSC. The dashed line indicates the minimum recommended threshold of 200,000 variants according to the LDSC documentation. Most GWAS retained approximately one million variants, and all retained more than 300,000 variants.

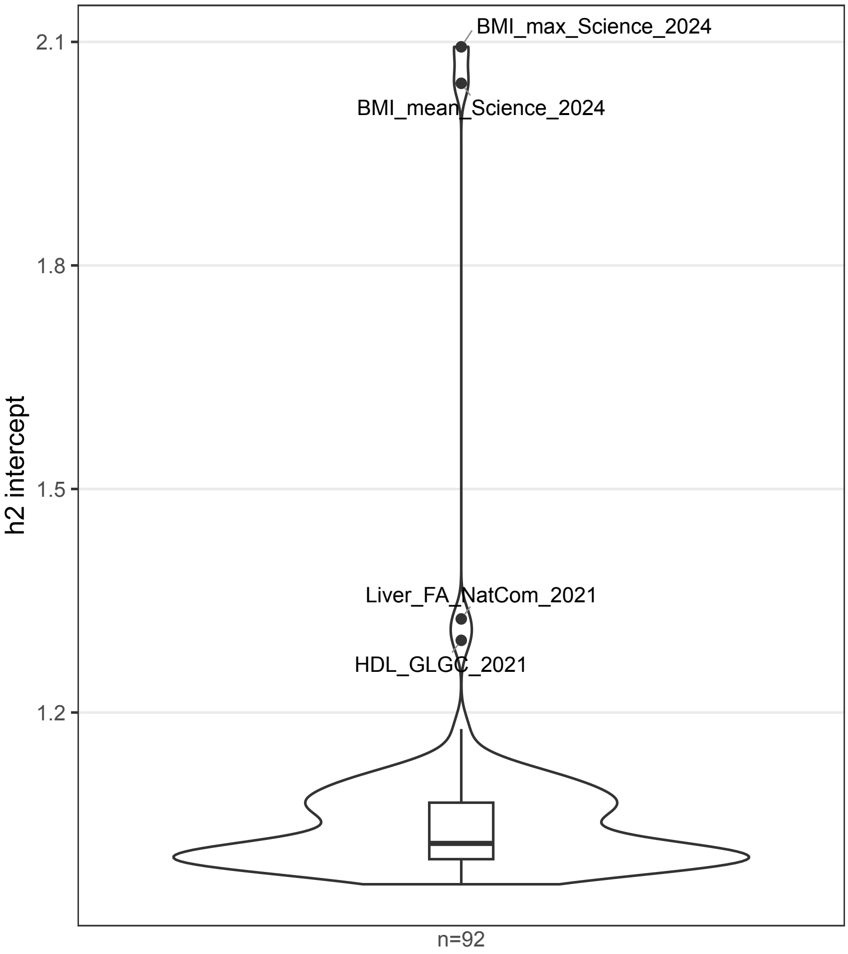


**Supplementary Fig. 6 | Distribution of LDSC intercept estimates across GWAS datasets.** Distribution of mean univariate LDSC intercept estimates across the 92 GWAS datasets with successful LDSC analyses, prior to quality-control filtering. For each trait, univariate intercept estimates obtained across pairwise LDSC analyses were averaged. Intercept values were inspected as indicators of residual inflation after accounting for polygenic signals.

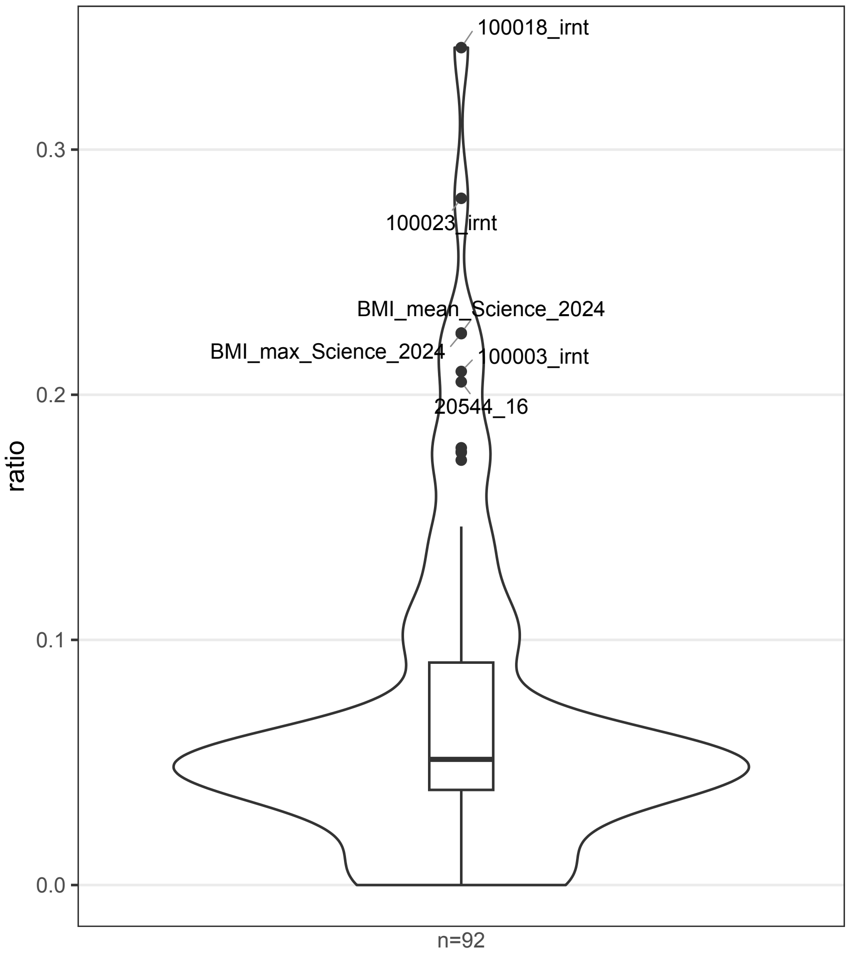


**Supplementary Fig. 7 | Distribution of LDSC attenuation ratios across GWAS datasets.** Distribution of mean univariate LDSC attenuation ratio estimates across the 92 GWAS datasets with successful LDSC analyses, prior to quality-control filtering. For each trait, univariate attenuation ratio estimates obtained across pairwise LDSC analyses were averaged. The attenuation ratio was used to assess the proportion of test-statistic inflation attributable to sources other than polygenic signal, with elevated values potentially indicating residual confounding or unreliable LDSC estimates. Values ≥0.20 were used to flag potentially problematic GWAS during quality-control assessment.

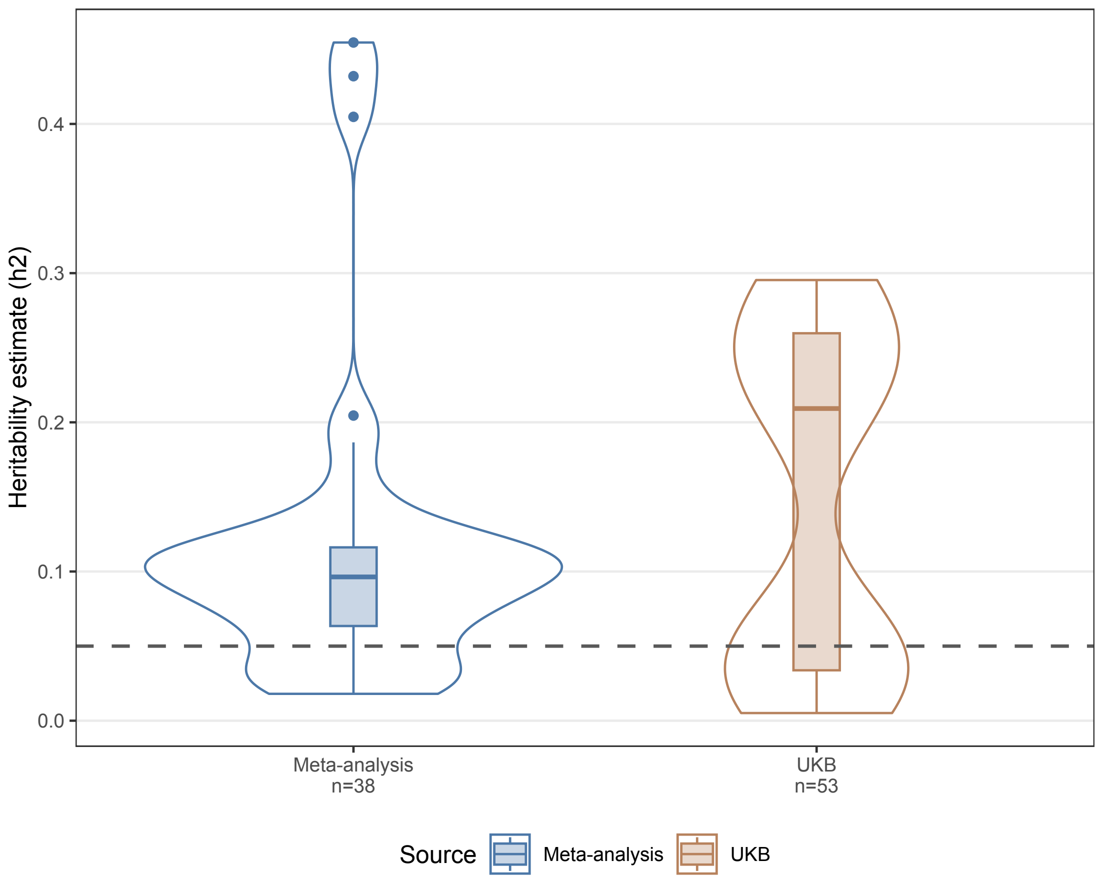


**Supplementary Fig. 8 | LDSC SNP-heritability estimates stratified by UK Biobank and non-UK Biobank studies.**

Distribution of mean univariate LDSC SNP-heritability estimates across the 92 GWAS datasets with successful LDSC analyses, prior to quality-control filtering, stratified according to whether they were primarily based on UK Biobank data or on non-UK Biobank cohorts and consortia. For each trait, univariate SNP-heritability estimates obtained across pairwise LDSC analyses were averaged. SNP-heritability estimates were used to assess the heritable signal available for downstream genetic-correlation analyses. The dashed line indicates the study-specific threshold of $h^2 = 0.05$, with traits excluded if any individual estimate fell below this threshold.

#### Supplementary Note 4. Diet-related traits clustering

This section provides additional information on the clustering strategies explored before defining the final diet-related trait clusters used in downstream analyses. We initially explored less computationally intensive approaches, applying PCA and k-means directly to GWAS summary statistics using either association *P* values or Z-scores across traits. These approaches were useful as exploratory analyses but had two main limitations. First, direct clustering of GWAS summary statistics did not incorporate LDSC-derived diagnostic metrics, such as intercept, ratio and SNP heritability, which were later used to evaluate GWAS calibration, potential confounding and the amount of phenotypic variance captured by common variants. Second, the resulting clustering patterns were dominated by study sources, particularly by UK Biobank-derived phenotypes, rather than by interpretable biological relationships among traits.

When PCA was applied to GWAS association statistics, UK Biobank traits accounted for most of the visible structure, whereas traits derived from meta-analyses or external consortia tended to cluster near the centre of the projection (Supplementary Figs. 9 and 10). Similarly, k-means clustering of the same summary-statistic representations produced partitions largely driven by UK Biobank membership. Gap statistic supported three clusters when using *P* values and four clusters when using Z-scores; however, in both cases, the inferred clusters mainly separated or fragmented UK Biobank traits, while most non-UK Biobank traits were grouped together (Supplementary Figs. 11–14).

Because these exploratory approaches were strongly influenced by study source and did not account for LDSC-derived measures of GWAS suitability, the final clustering strategy was based on pairwise genetic correlations estimated with LDSC bivariate regression. This approach summarizes shared genetic architecture between traits after accounting for LD structure and was therefore considered more appropriate for identifying biologically interpretable groups of diet-related traits. The distribution of LDSC bivariate regression *P* values is shown in Supplementary Fig. 15. Most pairwise genetic correlations showed statistically significant nominal P values (78.34% out of 1,653 unique pairs), and around 56.44% show strong statistical support as illustrated by the zoomed distribution relative to the Bonferroni-corrected threshold in Supplementary Fig. 16. The 358 nominally non-significant pairs generally had estimates close to zero (median absolute $r_g$ = 0.0274; 97.77% with $|r_g| < 0.10$), although non-significance alone does not establish the absence of genetic correlation. The final pairwise genetic correlation matrix used for trait clustering is shown in Supplementary Fig. 17

Absolute genetic correlations were transformed into dissimilarities defined as $1 - |r_g|$, treating strong positive and negative correlations as evidence of shared genetic architecture. These dissimilarities were approximated using a four-dimensional interval multidimensional scaling (MDS) configuration, fitted with the smacof R package using `mds(dist.matrix, type = "interval", init = "random", itmax = 100000, ndim = 4)`, with the random seed set to 9. Interval MDS fits configuration distances to a linear transformation of the original dissimilarities. The D1–D2 projection was used for clustering because visual inspection showed apparent and interpretable trait groupings in this plane. These coordinates were not treated as principal axes ordered by explained variance, projection choice prioritized interpretable trait groupings and accurate local genetic neighbourhoods. Genetic correlations are not guaranteed to satisfy the properties of Euclidean distances. MDS was therefore used to obtain an approximate Euclidean representation of these relationships, within which clustering delineated discrete groups along a continuum of shared genetic architecture.

To obtain a discrete trait partition, the D1–D2 coordinates were used as input to a genetic algorithm-based k-medoids clustering procedure. A schematic overview of the algorithm is shown in Supplementary Fig. 18. For each $k$ clusters, the GA initialized 400 candidate solutions, each consisting of $k$ distinct observed traits sampled without replacement as cluster representatives or medoids of the MDS space. Remaining traits were assigned to their nearest representative using Euclidean distances; exact assignment ties were resolved by random sampling. Candidates for which any representative had no assigned non-representative trait received a fitness penalty. Fitness was evaluated using the Dunn index, calculated as the minimum distance between representatives divided by the largest cluster-specific mean distance of assigned traits to their representative, excluding the representative itself. Under elitist reproduction, the highest-scoring parents were selected, with random selection among fitness ties. Crossover preserved representatives shared by both parents and randomly redistributed their remaining representatives between two offspring. The number of representatives to mutate was sampled from a binomial distribution with size $k$ and probability 0.2; selected positions were replaced by distinct traits not already present in the candidate. Parents and offspring were pooled, and the 400 highest-scoring candidates were retained for the next generation. The procedure was run for 100 generations for $k$ = 2–10. The highest observed final fitness was obtained at $k$ = 4 (Supplementary Fig. 19), and fitness trajectories are shown in Supplementary Fig. 20.

Following clustering, we reviewed extreme intercluster genetic associations, defined as absolute genetic correlations ≥ 0.8, to identify traits whose assignment might separate strongly shared genetic architectures. Such associations involved Weight and body-fat traits, Magnesium and Food weight, and DHA and Omega3. For each implicated trait, we compared the representation of its original cluster and the candidate neighbouring cluster among its ten strongest absolute genetic correlations. Reassignment required both an extreme external association and greater representation of the candidate cluster than the original cluster. DHA was the only trait meeting both criteria: four of its ten strongest associations involved C3, compared with two involving its original cluster, C4. It was therefore reassigned to C3 for downstream analyses. Weight showed equal representation of C2 and C4 among its ten strongest associations (five each), while the other implicated traits retained comparable or greater support within their original clusters and were not reassigned. The Voronoi boundaries in Fig. 1 retain the original GA partition.

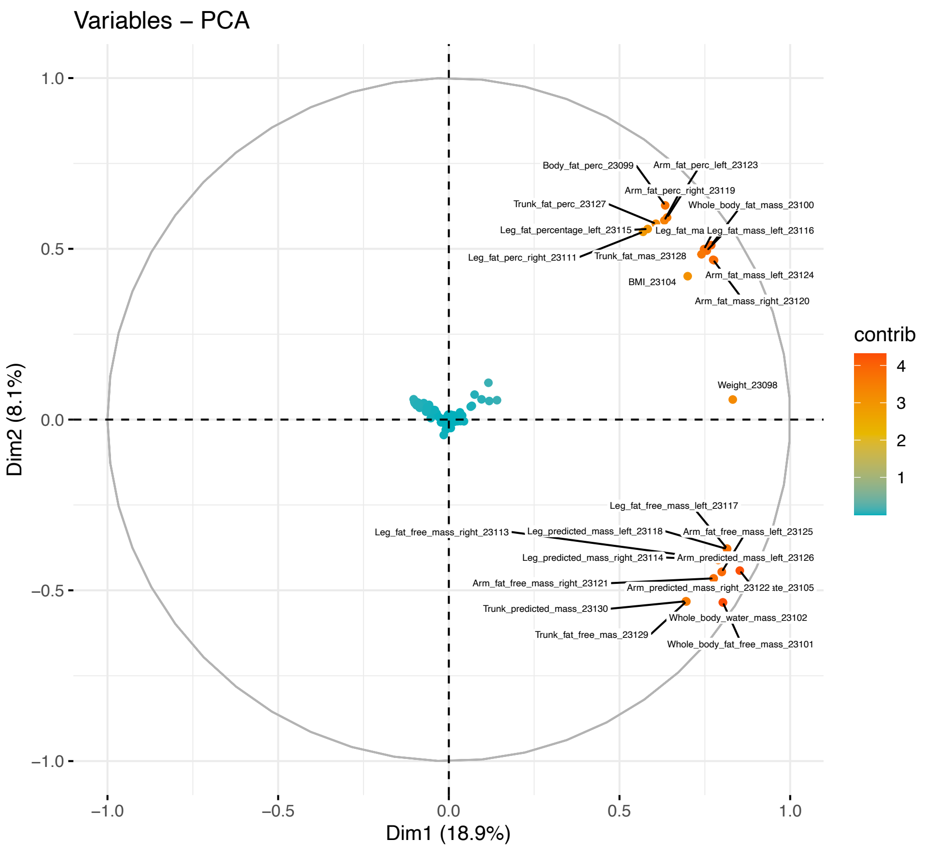


**Supplementary Fig. 9 | PCA of GWAS association P values across diet-related traits.** Principal component analysis was applied directly to GWAS association P values across the initially considered diet-related traits. UK Biobank-derived traits drive most of the visible structure, whereas traits from external meta-analyses and consortia tend to project near the centre of the PCA space, indicating that direct clustering of GWAS summary statistics was strongly influenced by study source.

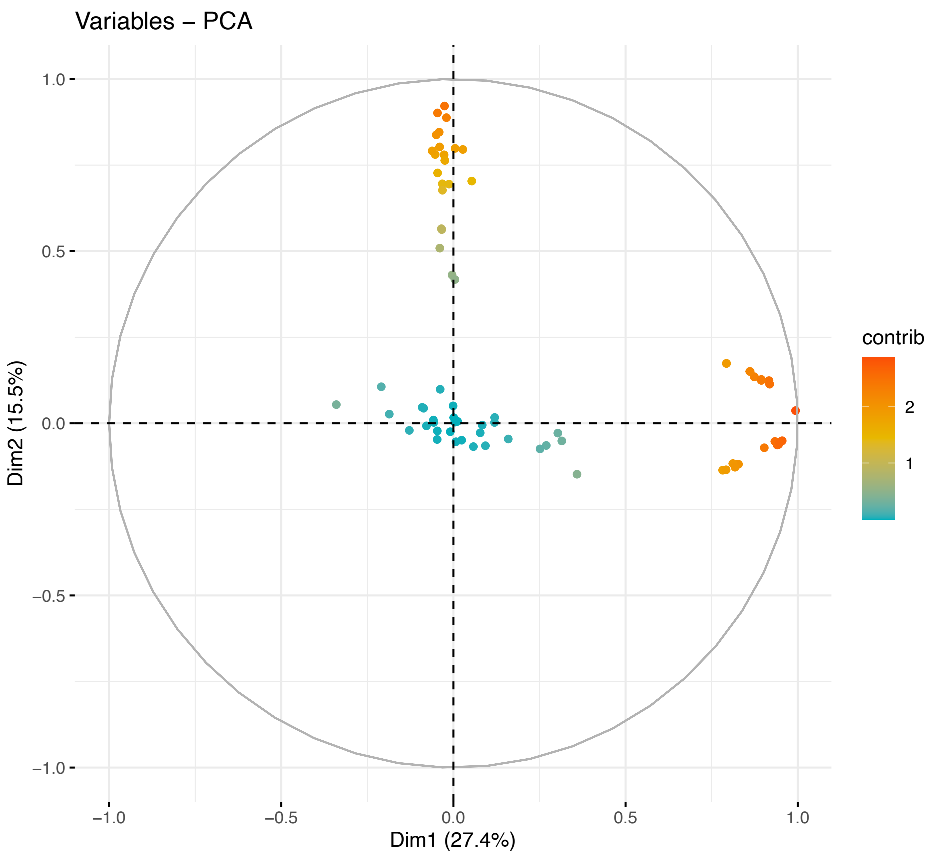


**Supplementary Fig. 10 | PCA of GWAS Z-scores across diet-related traits.** Principal component analysis was applied directly to GWAS Z-scores across diet-related traits. As observed with P values, UK Biobank-derived traits account for most of the visible clustering structure, while non-UK Biobank traits show limited separation in the projected space.

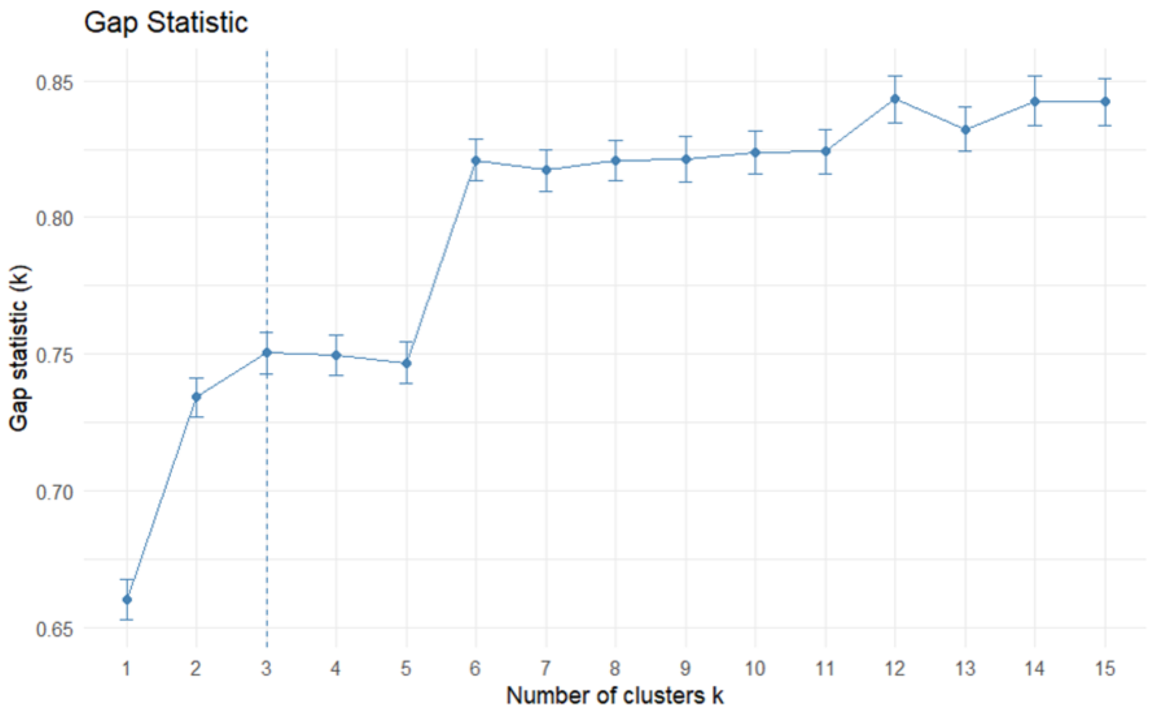


**Supplementary Fig. 11 | Gap statistic for k-means clustering using GWAS association P values.**

Gap statistic analysis was used to evaluate the optimal number of k-means clusters when clustering traits directly from GWAS association P values. The optimal solution supported three clusters, which largely reflected separation of UK Biobank-derived traits from non-UK Biobank traits rather than biologically interpretable diet-related domains.

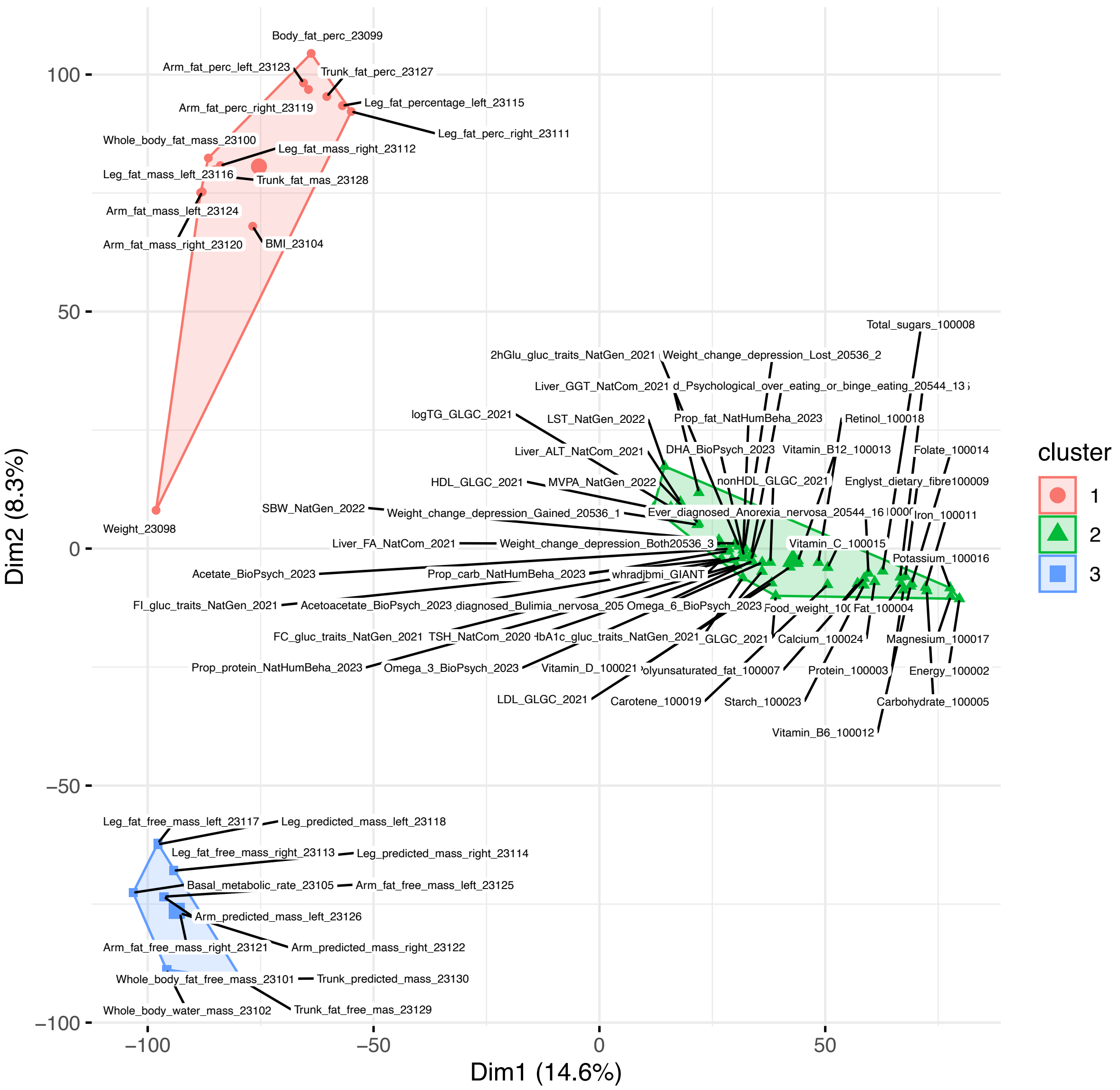


**Supplementary Fig. 12 | K-means clustering of diet-related traits using GWAS association P values.**
K-means clustering was applied to GWAS association P values using the optimal number of clusters inferred from the gap statistic. The resulting partition was dominated by UK Biobank-derived traits, while most traits from external meta-analyses and consortia were grouped together.

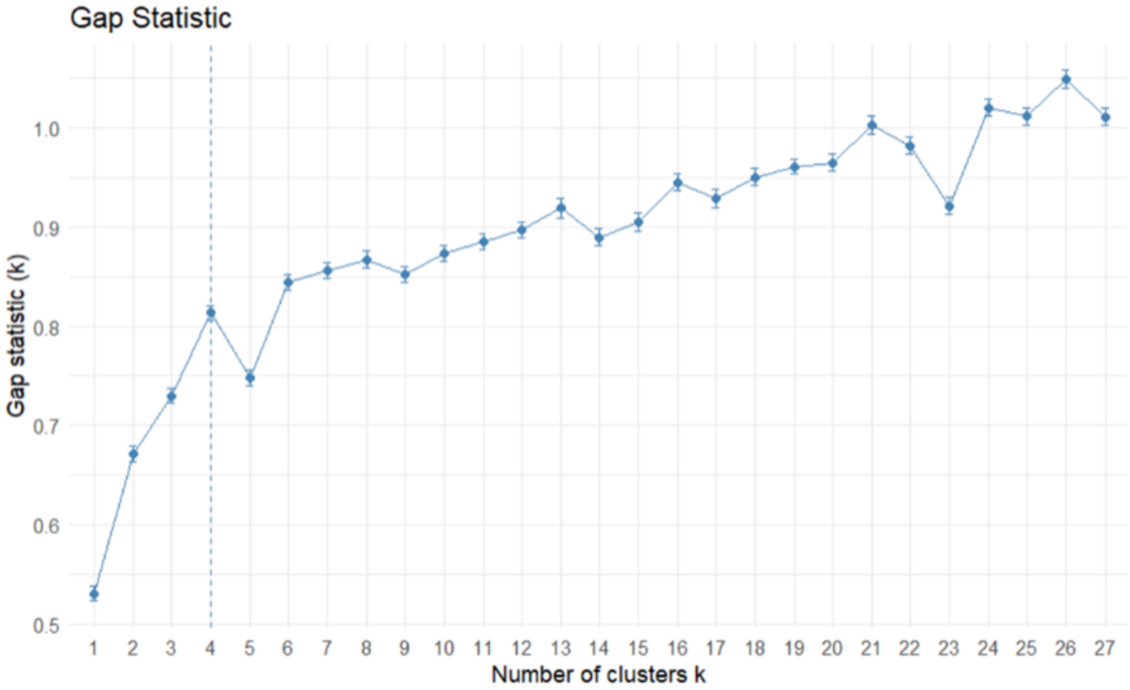


**Supplementary Fig. 13 | Gap statistic for k-means clustering using GWAS Z-scores.**
Gap statistic analysis was used to evaluate the optimal number of k-means clusters when clustering traits directly from GWAS Z-scores. The optimal solution supported four clusters, but the inferred structure remained largely driven by UK Biobank-derived traits.

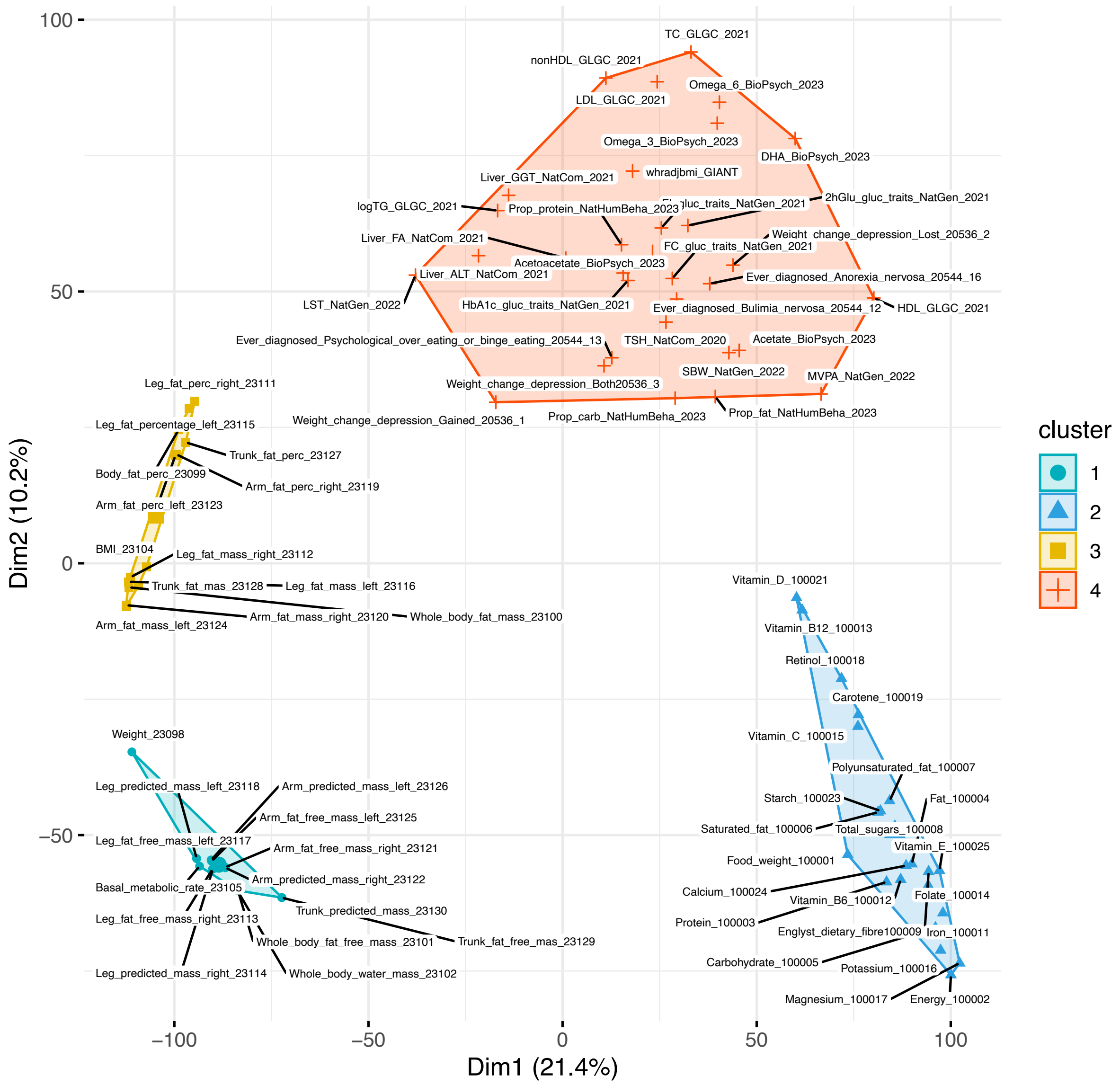


**Supplementary Fig. 14 | K-means clustering of diet-related traits using GWAS Z-scores.**

K-means clustering was applied to GWAS Z-scores using the optimal number of clusters inferred from the gap statistic. Compared with the P-value-based clustering, UK Biobank traits showed greater fragmentation across clusters, while non-UK Biobank traits remained weakly differentiated.

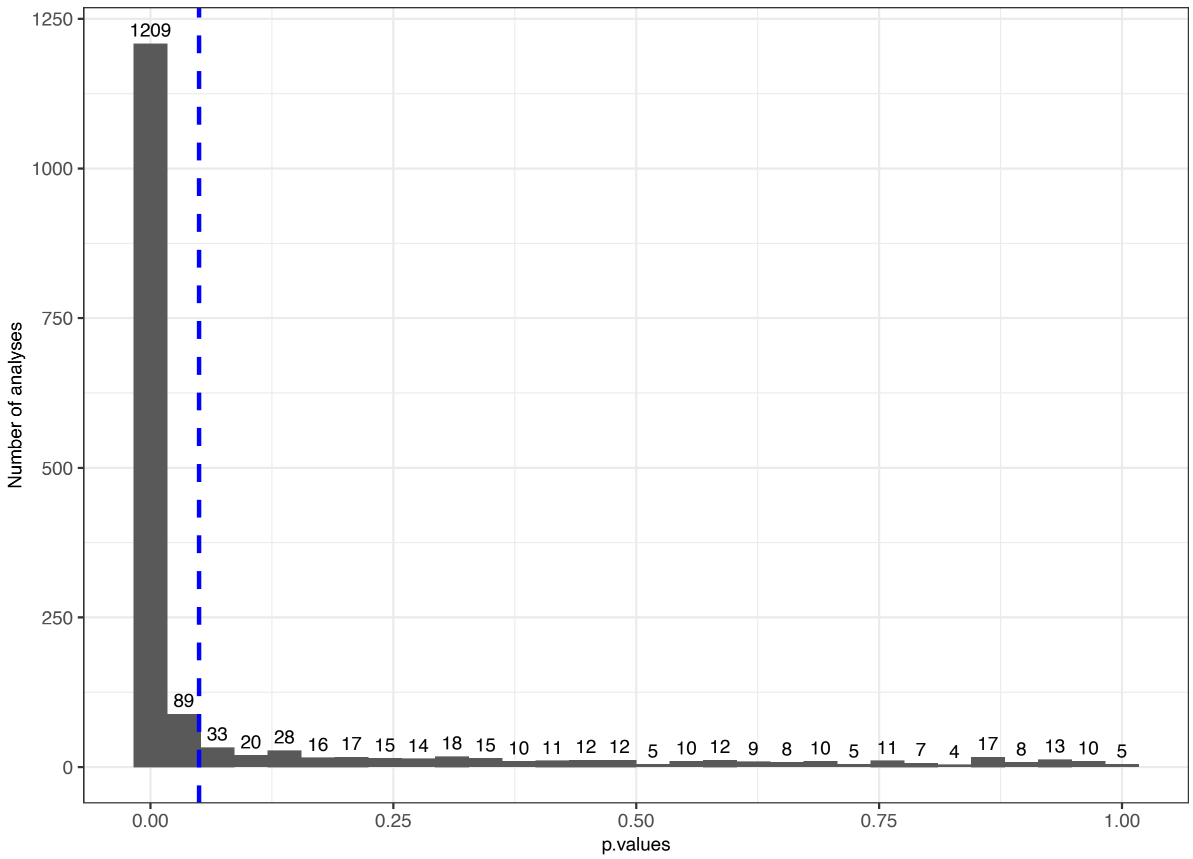


**Supplementary Fig. 15 | Distribution of LDSC bivariate regression P values across trait pairs.**

Distribution of P values obtained from pairwise LDSC bivariate regressions across the final set of 58 diet-related traits. The dashed line indicates the nominal significance threshold of P = 0.05.

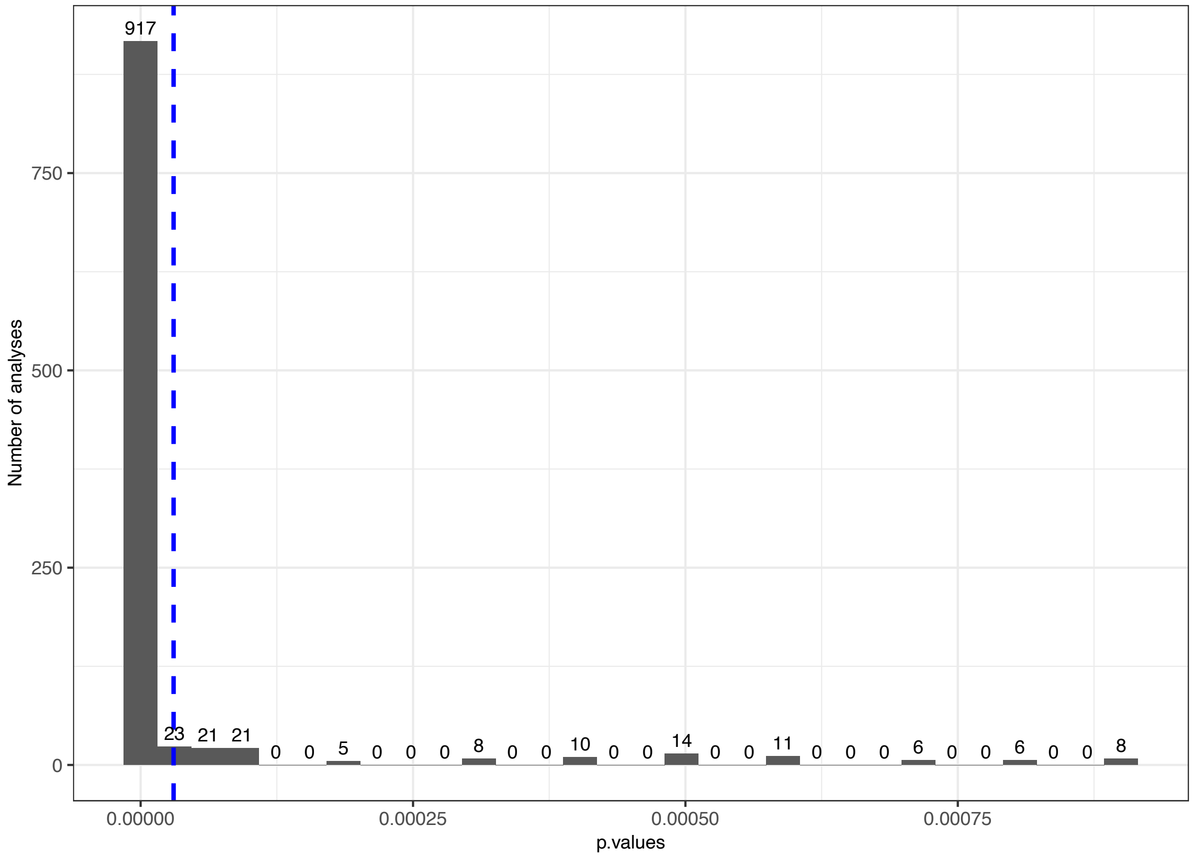


**Supplementary Fig. 16 | Zoomed distribution of LDSC bivariate regression P values relative to Bonferroni correction.**

Zoomed view of the lower tail of the LDSC bivariate regression P-value distribution across pairwise genetic correlations. The dashed line indicates the Bonferroni-corrected threshold for 1,653 pairwise tests ($P \approx 3.0 \times 10^{-5}$).

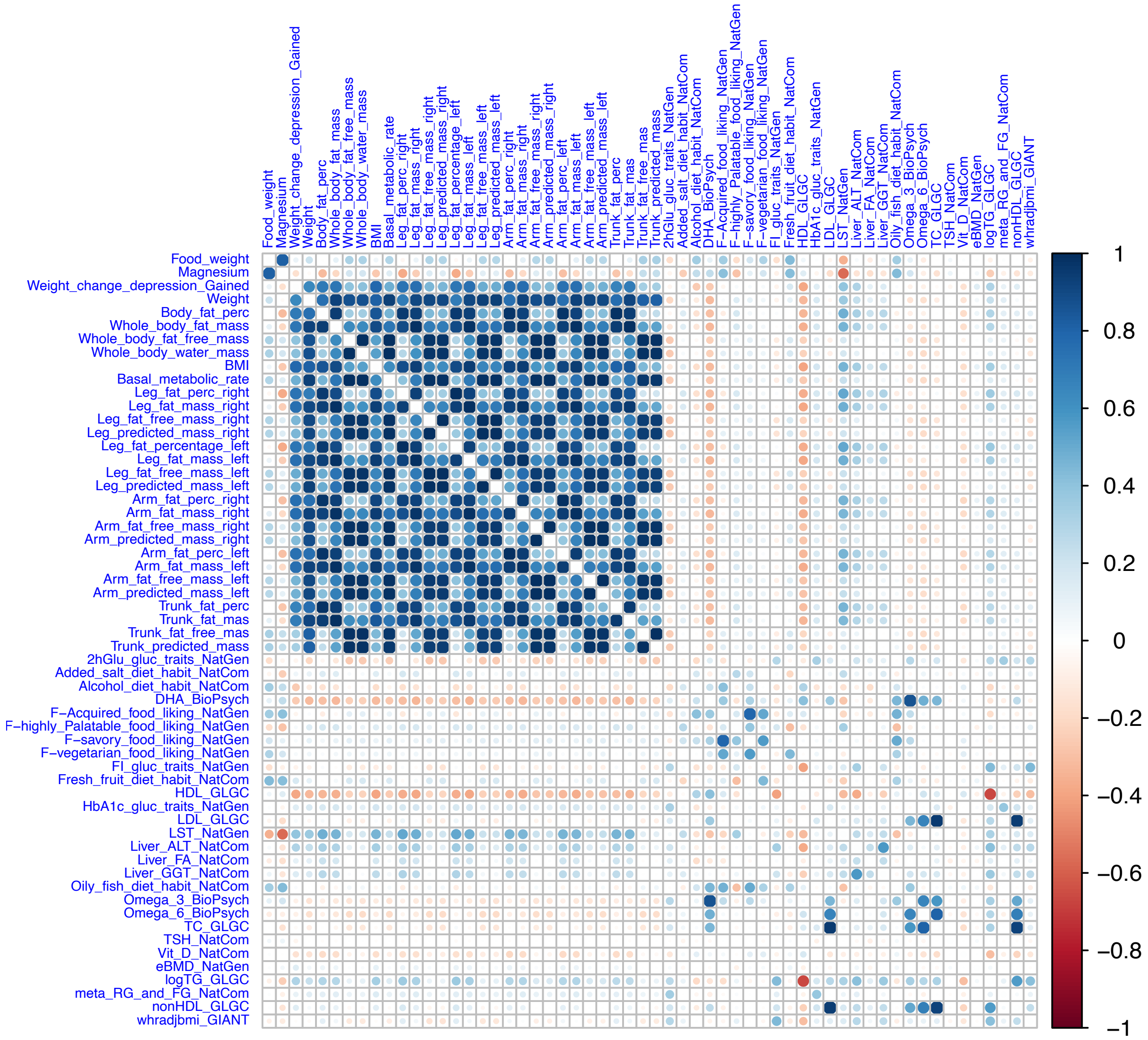


**Supplementary Fig. 17 | Pairwise genetic correlation matrix among the 58 diet-related traits.**

Matrix of pairwise genetic correlations estimated using LDSC bivariate regression for the final set of 58 diet-related traits retained after quality control. Colours represent the direction and magnitude of genetic correlation coefficients, with positive and negative values indicating agonistic and antagonistic shared genetic architecture, respectively.

**Algorithm 1: Genetic algorithm for medoid-based clustering in MDS space**

**Input:** Trait labels $D = \{d_1, \ldots, d_n\}$; MDS coordinates D1 and D2 for each trait; even population size $P$; number of clusters $k$; mutation rate $r \in [0, 1]$; generations $G$
**Output:** Best fitness $f_{\text{best}}$ and best medoid solution $M_{\text{best}}$
`// Euclidean distance in the two-dimensional MDS representation`
$\delta(u, v) = \sqrt{(\mathrm{D1}_u - \mathrm{D1}_v)^2 + (\mathrm{D2}_u - \mathrm{D2}_v)^2}$ for any traits $u, v \in D$
**Function** `Evaluate(`$M$`)`:
- Let $M = (m_1, \ldots, m_k)$ be the selected medoid labels
- Initialize empty clusters $C_1, \ldots, C_k$
- `//` $C_j$ `contains the traits assigned to medoid` $m_j$
- **foreach** $d \in D \setminus M$ **do**
  - $j^* \leftarrow \arg\min_{1 \le j \le k} \delta(d, m_j)$ `// Resolve ties at random`
  - Add trait $d$ to $C_{j^*}$
- **if** *any* $C_j$ *is empty* **then**
  - **return** $-100$ `// Extreme penalty to discourage retention in the population`
- `// Mean within-cluster distance to the medoid`
- $\bar{\delta}_j \leftarrow \mathrm{mean}_{d \in C_j} \delta(d, m_j)$ for $j = 1, \ldots, k$
- `// Dunn-type fitness: minimum medoid separation / maximum within-cluster mean`
- **return** $f(M) = \dfrac{\min_{1 \le a < b \le k} \delta(m_a, m_b)}{\max_{1 \le j \le k} \bar{\delta}_j}$
- `//` $a < b$ `considers each pair of distinct medoids once; maximize` $f(M)$

`// Initialization`
$S \leftarrow (M_1, \ldots, M_P)$, with each $M_i \leftarrow$ `SampleWithoutReplacement(`$D, k$`)`
**foreach** $M_i \in S$ *in parallel* **do**
- $f_i \leftarrow$ `Evaluate(`$M_i$`)`

**for** $g \leftarrow 1$ **to** $G$ **do**
- $O \leftarrow \varnothing$
- **for** $i \leftarrow 1$ **to** $P/2$ in parallel **do**
  - `// Elitist parent selection, repeated for every mating`
  - Select distinct population entries $p_1, p_2$ with the two highest fitness values; sample uniformly among tied candidates
  - `// Recombination: preserve common medoids in both children`
  - $U \leftarrow p_1 \cap p_2$; $V \leftarrow (p_1 \setminus U) \cup (p_2 \setminus U)$
  - $(V_1, V_2) \leftarrow$ `RandomEqualPartition(`$V$`)`
  - $c_1 \leftarrow U \cup V_1$; $c_2 \leftarrow U \cup V_2$
  - **foreach** $c \in \{c_1, c_2\}$ **do**
    - `// Mutation: draw replacements from outside the pre-mutation child`
    - $B \sim \mathrm{Binomial}(k, r)$
    - Select $B$ positions uniformly without replacement
    - Replace them with $B$ distinct labels sampled from $D \setminus c$
  - Append $c_1, c_2$ to $O$
- **foreach** $M_i \in O$ *in parallel* **do**
  - $f_i \leftarrow$ `Evaluate(`$M_i$`)`
- `// Elitist population update; retain multiplicities`
- $S \leftarrow$ the $P$ highest-fitness entries of the concatenation $(S, O)$

$(f_{\text{best}}, M_{\text{best}}) \leftarrow$ highest stored fitness and corresponding solution in $S$
**return** $(f_{\text{best}}, M_{\text{best}})$

**Supplementary Fig. 18 | Schematic overview of the genetic algorithm-based clustering procedure.**

Workflow of the genetic algorithm used to optimize clustering of diet-related traits in the two-dimensional MDS space derived from pairwise genetic correlations. For a given number of clusters ($k$), the algorithm starts by generating an initial population of candidate solutions, each defined by a set of observed traits chosen as medoids. Traits are assigned to the nearest medoid according to their MDS coordinates. The fitness of each solution is evaluated using the Dunn Index, which quantifies clustering quality by maximizing inter-cluster separation while minimizing intra-cluster compactness. The best-performing solutions are retained and used to generate a new population through crossover and mutation. Mutation sites are sampled from a binomial distribution using the specified mutation rate, allowing random replacement of medoids while preserving diversity in the search space. This iterative procedure is repeated for 100 generations and for values of $k$ ranging from 2 to 10 in order to identify the optimal clustering solution.

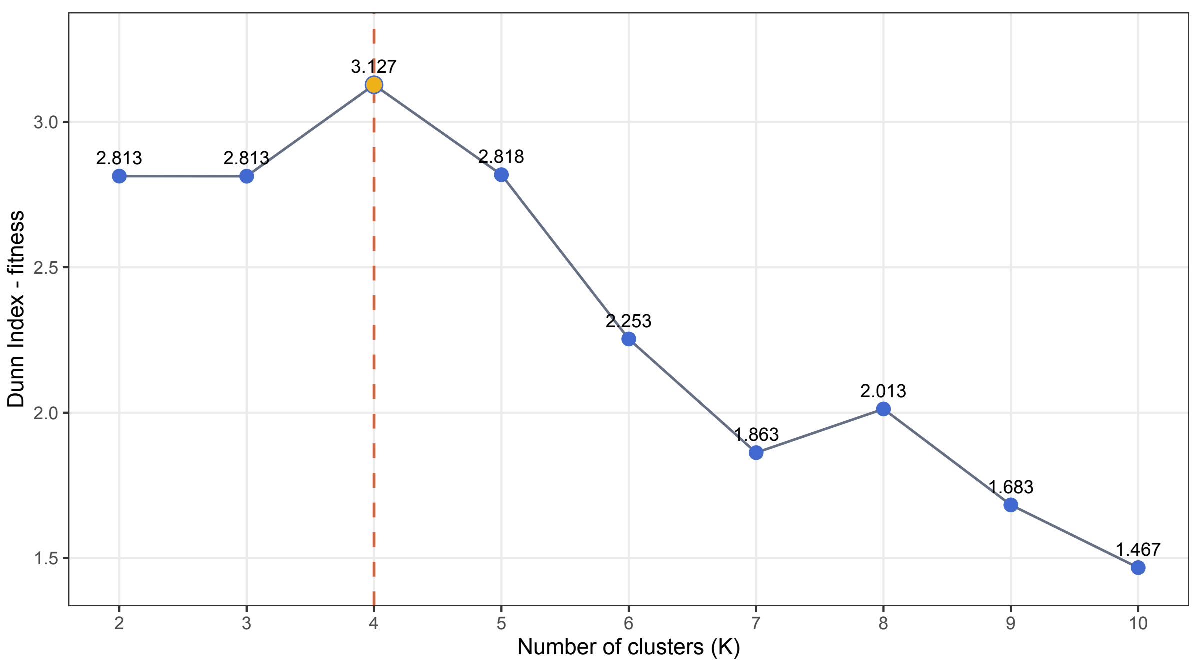


**Supplementary Fig. 19 | Dunn Index across candidate numbers of trait clusters.** Dunn Index values obtained for clustering solutions with $k$ ranging from 2 to 10. Higher values indicate better clustering quality, corresponding to greater inter-cluster separation and lower intra-cluster compactness. The dashed line marks the maximum Dunn Index, which was observed for $k$ = 4 and selected as the optimal clustering solution.

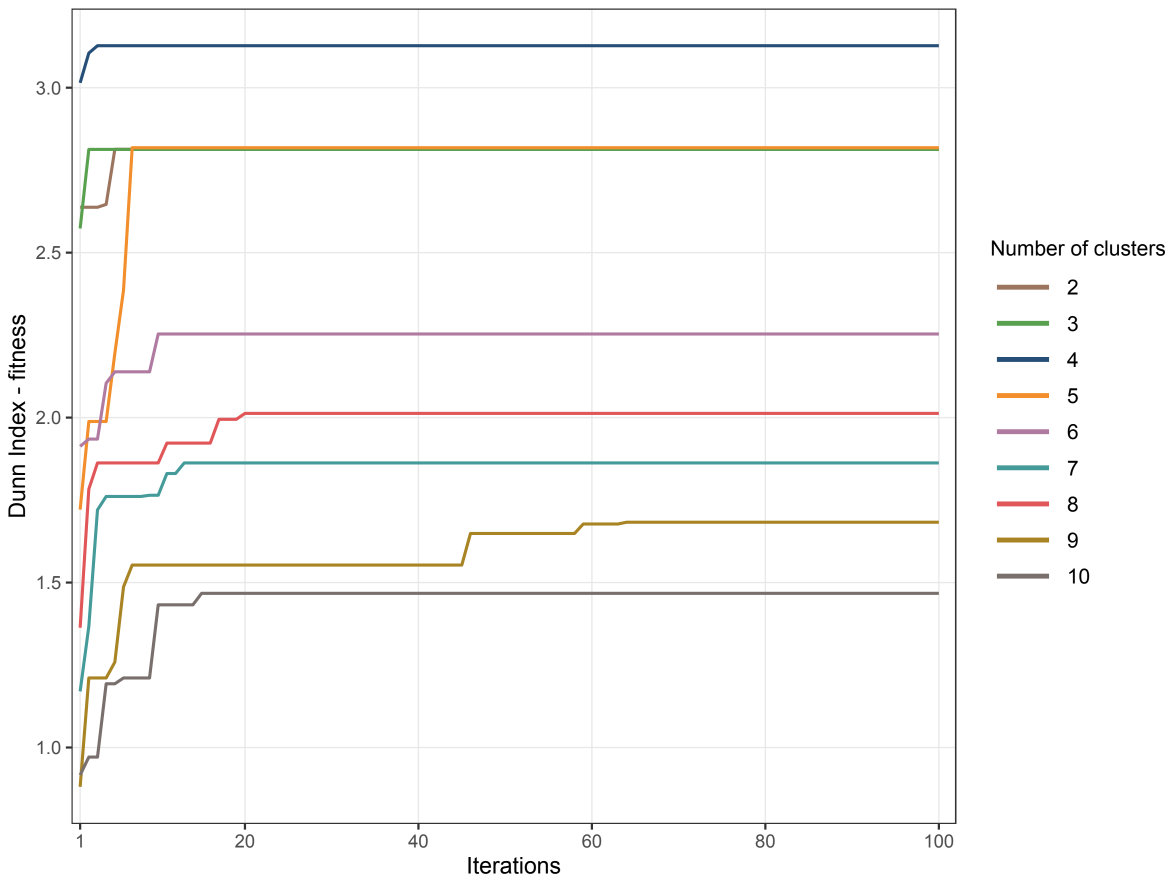


**Supplementary Fig. 20 | Convergence of the genetic algorithm across candidate numbers of clusters.**

Fitness trajectories of the genetic algorithm across 100 iterations for clustering solutions with *k* ranging from 2 to 10. The y-axis represents the fitness value, defined by the Dunn Index. These trajectories illustrate the convergence behaviour of the algorithm and the iteration at which the best solution was reached for each value of *k*.

#### Supplementary Note 5. Temporal association testing and null models

This section provides additional visual summaries of the temporal association framework and the null models used to evaluate whether the observed trajectories of diet-related traits differed from random expectation. Supplementary Fig. 21 summarizes the cluster-level permutation test used to assess whether the number of significant traits observed after within-cluster Benjamini–Hochberg correction exceeded random expectation in cluster 1. Supplementary Figs. 22–25 show the temporal trajectories of all tested traits grouped by genetic cluster, including weighted GAM smooths for visualization and the corresponding weighted Kendall's $\tau$ and $P$ values. For these plots, sample age is represented in years before present (BP); therefore, positive weighted Kendall's $\tau$ values indicate increasing risk-allele proportions towards the present, whereas negative values indicate decreasing proportions towards the present. Supplementary Fig. 26 summarizes the residual genetic correlation structure among traits that remained significant after multiple-testing correction. An example null distribution of weighted Kendall's $\tau$ derived from random risk-allele permutation is shown in Supplementary Fig. 27, if retained.

Temporal associations were tested using weighted Kendall's $\tau$, with empirical significance assessed by 10,000 randomizations of risk-allele assignments, as described in Methods. To evaluate whether the concentration of significant traits within a given genetic cluster could arise by chance, empirical $P$ values were permuted across clusters and Benjamini–Hochberg correction was recalculated at each iteration, generating a null distribution for the number of significant traits expected per cluster. This framework allowed observed cluster-level enrichment to be compared against random expectation while preserving the multiple-testing structure of the analysis.

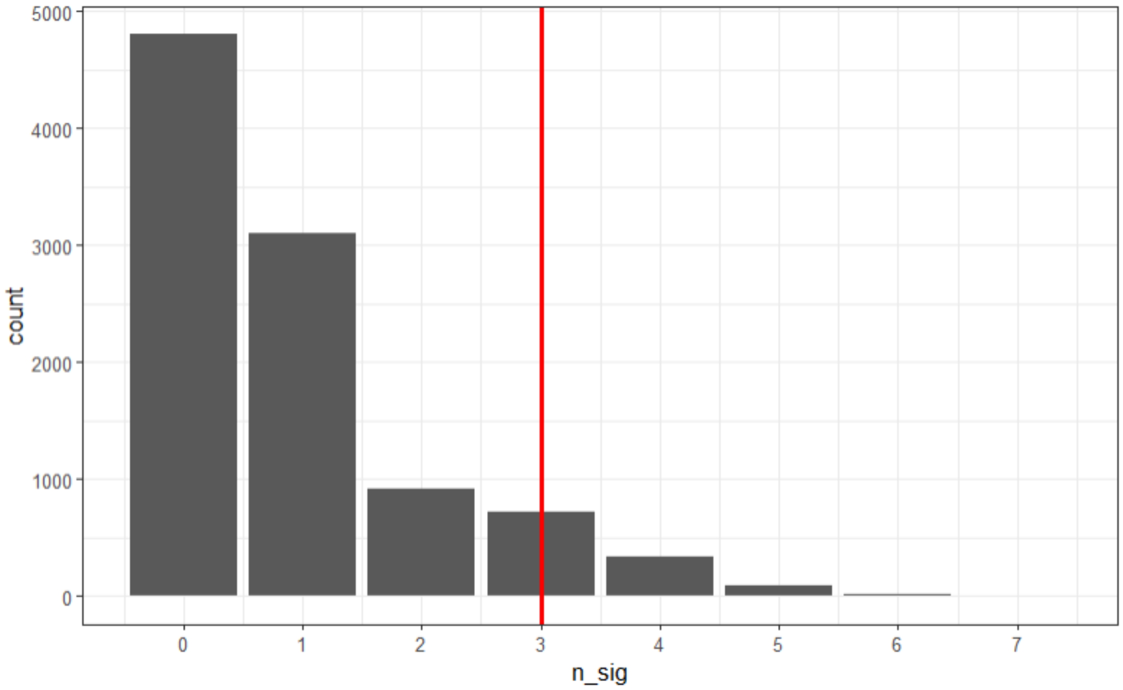


**Supplementary Fig. 21 | Null distribution of the number of significant traits after within-cluster BH correction.**

Discrete null distribution obtained by permuting empirical $P$ values across the four genetic clusters and recalculating the number of significant traits after Benjamini–Hochberg correction within each cluster across 10,000 iterations. The x-axis indicates the number of significant traits after correction, and the y-axis indicates the number of permutations yielding each outcome. The vertical red line marks the observed number of significant traits in cluster 1. This analysis was used to evaluate whether the apparent enrichment of significant temporal signals within cluster 1 differed from random expectation.

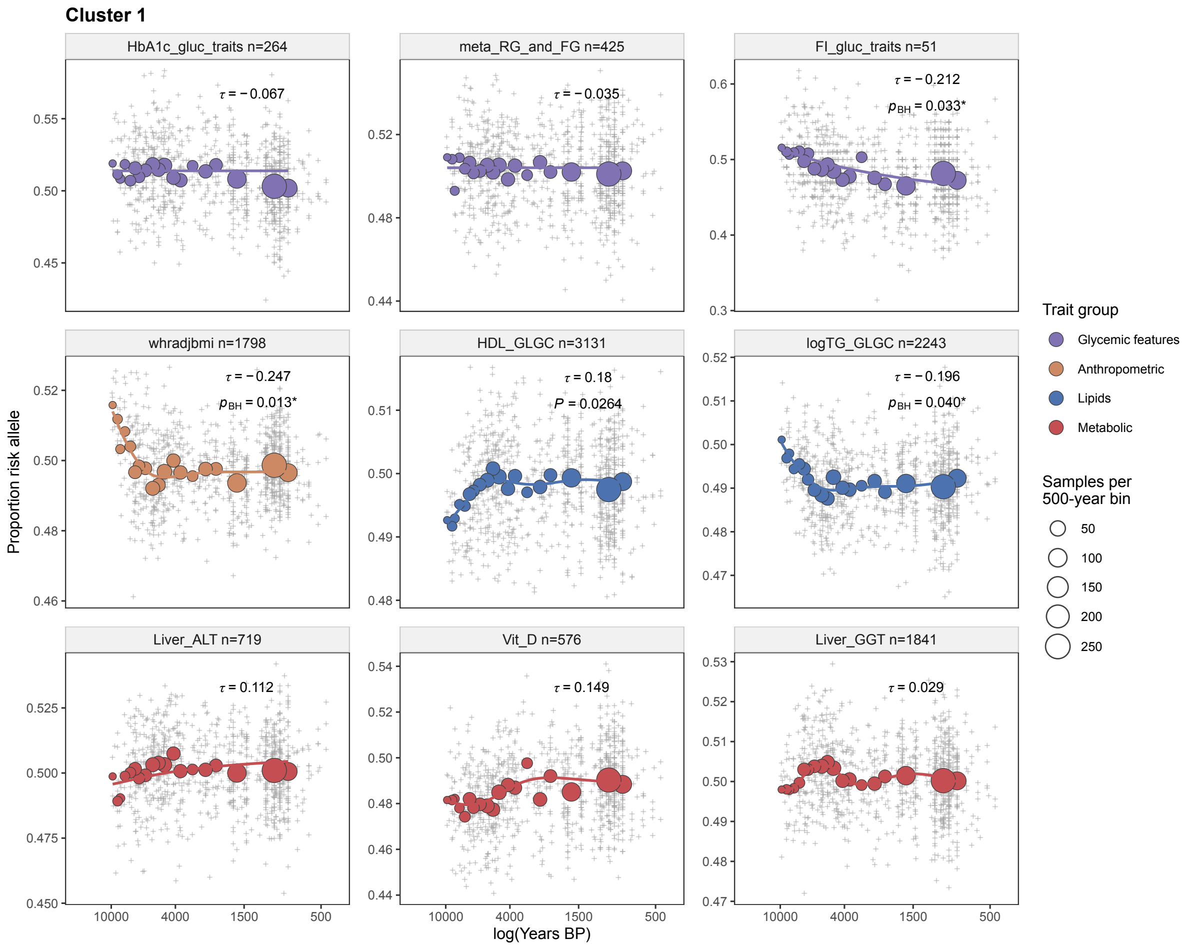


**Supplementary Fig. 22 | Temporal trajectories of diet-related traits in cluster 1.** Temporal trajectories for traits assigned to cluster 1, showing the proportion of risk alleles across ancient individuals over time. The x-axis represents sample age in years before present (BP) on a log scale, and the y-axis represents the mean proportion of risk alleles. A weighted GAM smooth is shown for visualization, using the same temporal weights applied in the weighted Kendall's $\tau$ analyses. Each panel reports the corresponding weighted Kendall's $\tau$ statistic and the associated empirical $P$ value or Benjamini–Hochberg-corrected $P_{BH}$ value when applicable.

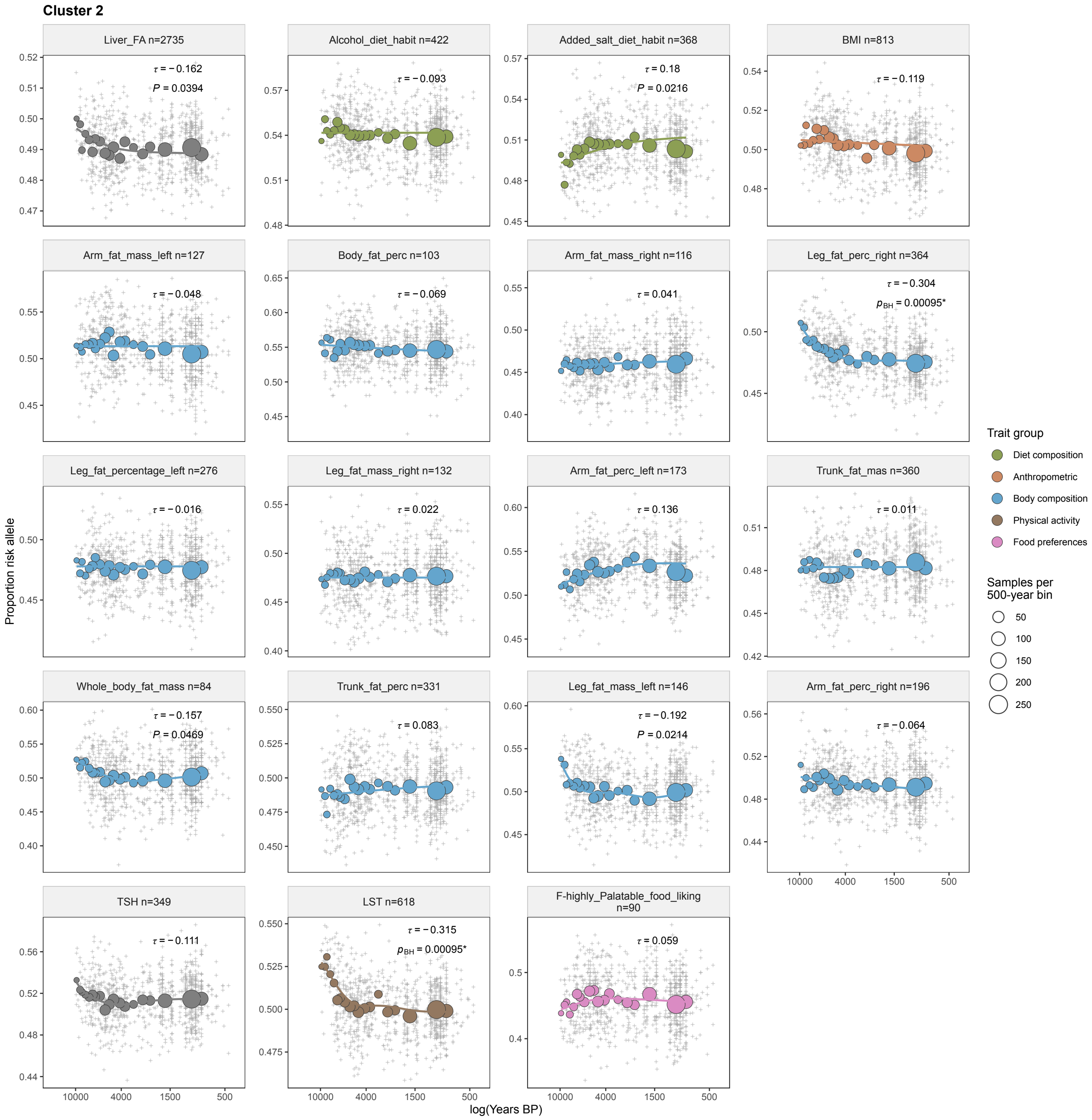


**Supplementary Fig. 23 | Temporal trajectories of diet-related traits in cluster 2.**
Temporal trajectories for traits assigned to cluster 2, showing the proportion of risk alleles across ancient individuals over time. The x-axis represents sample age in years before present (BP) on a log scale, and the y-axis represents the mean proportion of risk alleles. A weighted GAM smooth is shown for visualization, using the same temporal weights applied in the weighted Kendall's $\tau$ analyses. Each panel reports the corresponding weighted Kendall's $\tau$ statistic and the associated empirical $P$ value or Benjamini–Hochberg-corrected $P_{BH}$ value when applicable.

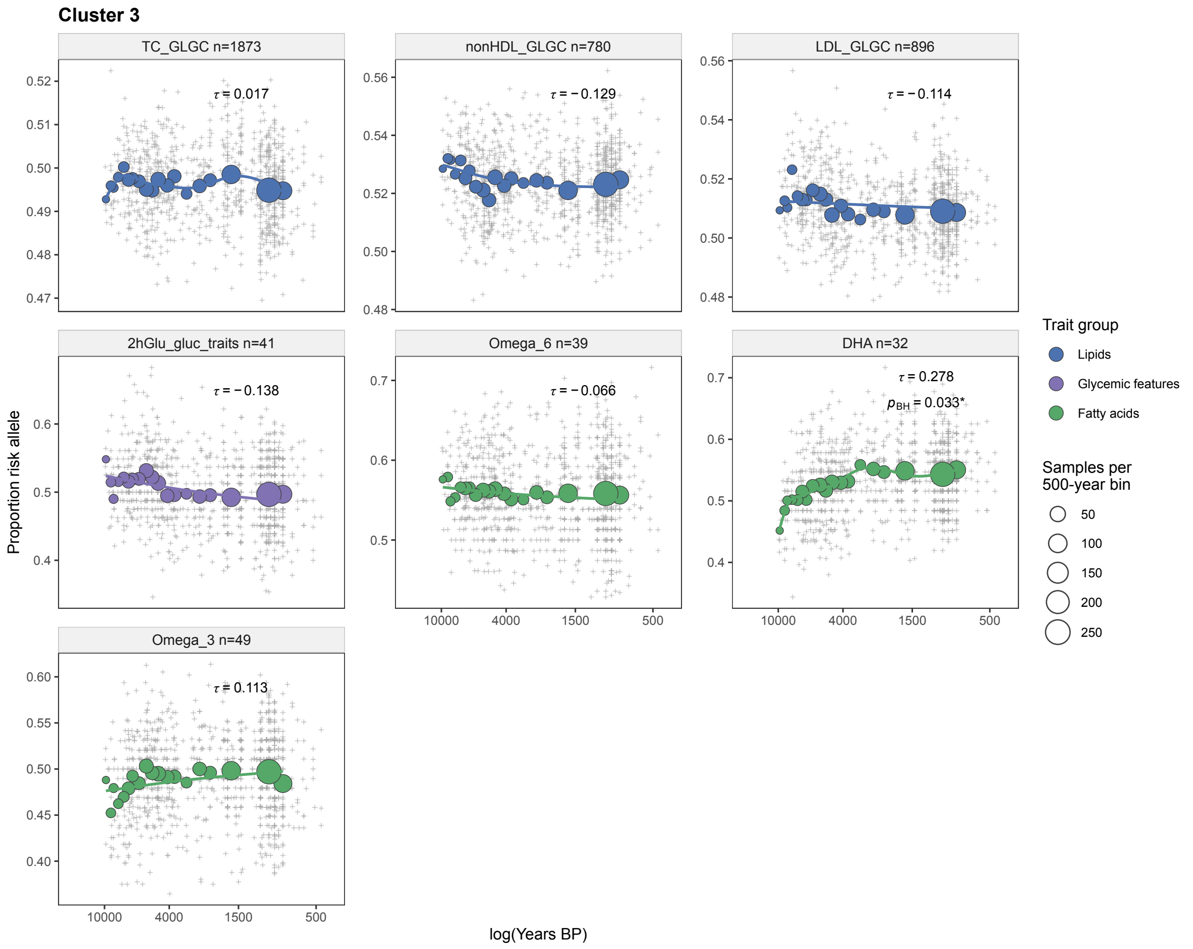


**Supplementary Fig. 24 | Temporal trajectories of diet-related traits in cluster 3.**
Temporal trajectories for traits assigned to cluster 3, showing the proportion of risk alleles across ancient individuals over time. The x-axis represents sample age in years before present (BP) on a log scale, and the y-axis represents the mean proportion of risk alleles. A weighted GAM smooth is shown for visualization, using the same temporal weights applied in the weighted Kendall's $\tau$ analyses. Each panel reports the corresponding weighted Kendall's $\tau$ statistic and the associated empirical $P$ value or Benjamini–Hochberg-corrected $P_{BH}$ value when applicable.

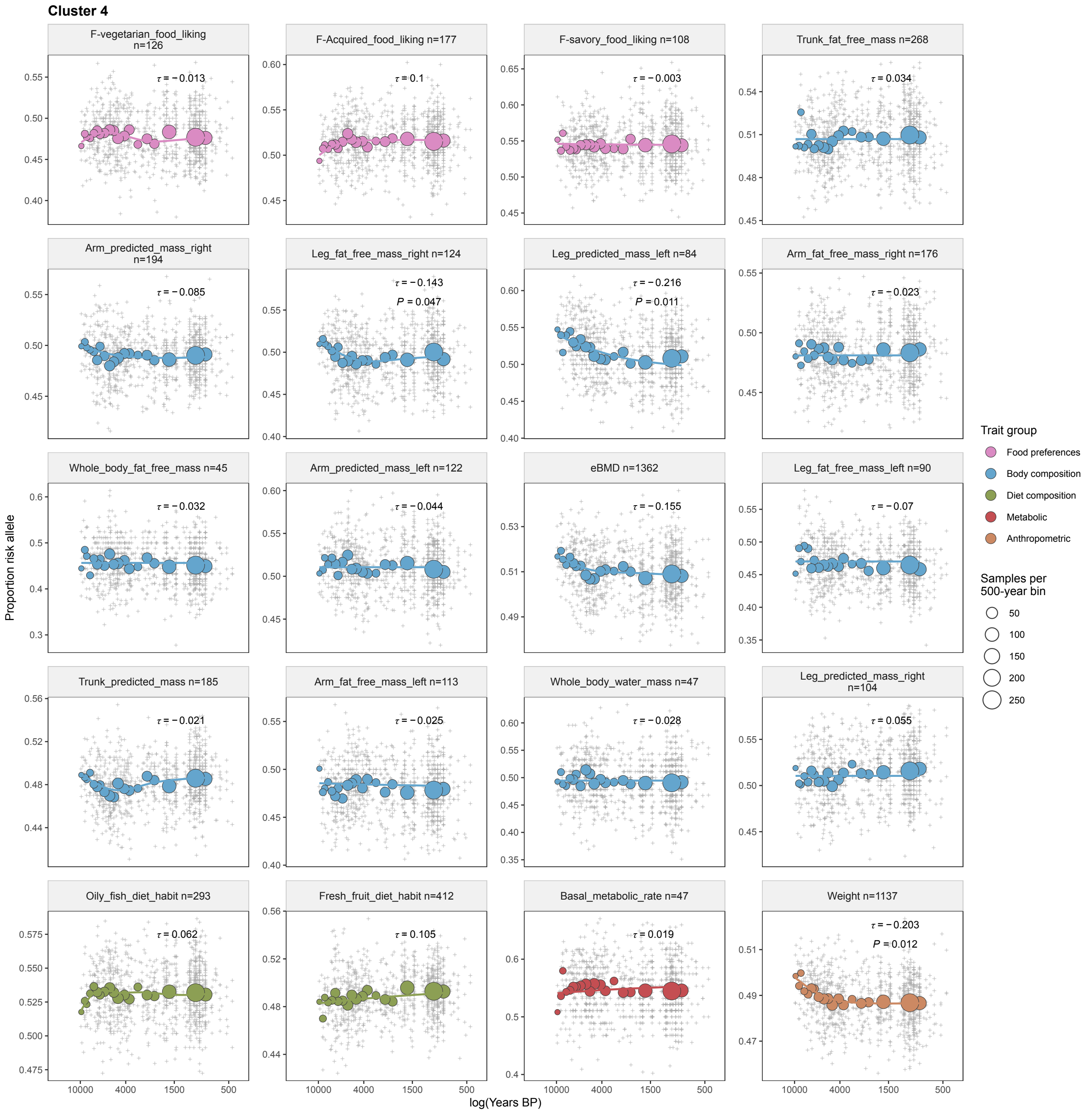


**Supplementary Fig. 25 | Temporal trajectories of diet-related traits in cluster 4.** Temporal trajectories for traits assigned to cluster 4, showing the proportion of risk alleles across ancient individuals over time. The x-axis represents sample age in years before present (BP) on a log scale, and the y-axis represents the mean proportion of risk alleles. A weighted GAM smooth is shown for visualization, using the same temporal weights applied in the weighted Kendall's $\tau$ analyses. Each panel reports the corresponding weighted Kendall's $\tau$ statistic and the associated empirical $P$ value or Benjamini–Hochberg-corrected $P_{BH}$ value when applicable.

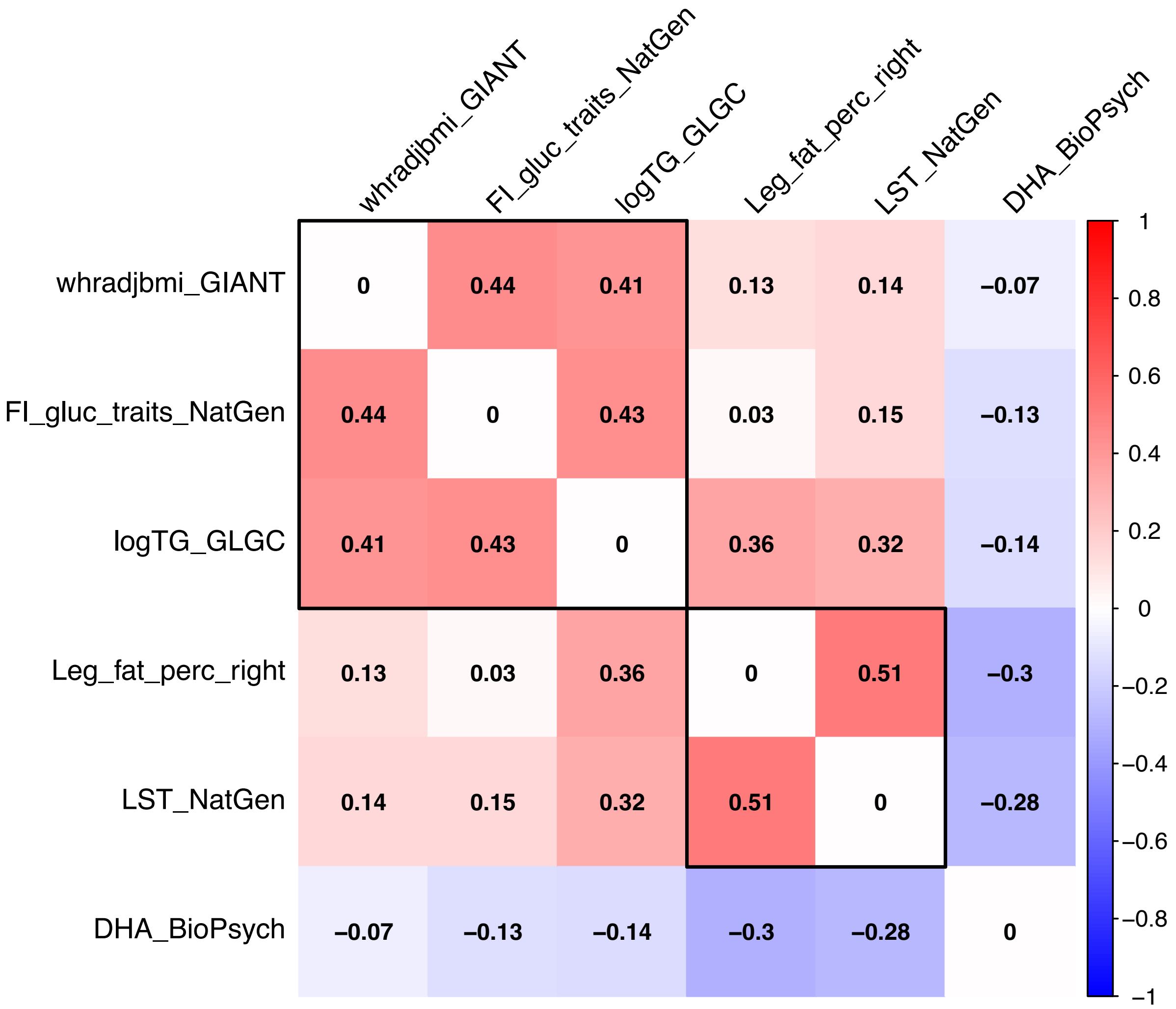


**Supplementary Fig. 26 | Pairwise genetic correlation matrix among traits significant after BH correction.**

Pairwise LDSC genetic correlations among traits that remained significant after within-cluster Benjamini–Hochberg correction. The matrix illustrates that traits within the same cluster tend to retain moderate to strong correlations, whereas residual correlations between significant traits from different clusters are comparatively weaker. Colours represent the direction and magnitude of the genetic correlation coefficients, and black boxes delimitate traits belonging to the same cluster.

#### Supplementary Note 6. Empirical assessment against a genome-wide background sensitive to demographic history and background selection

This section provides additional details and visual summaries of the empirical assessment used to evaluate whether the temporal trajectories that remained significant after within-cluster FDR correction exceeded the variation observed in a genome-wide background potentially sensitive to demographic history and background selection. The null model was constructed from a genome-wide set of 1,844,650 markers linked to 13,172 protein-coding genes, including variants located within genes and within 50 kb of gene boundaries. For each tested trait, 1,000 proxy marker sets were sampled by matching allele frequency and marker number to the observed trait-specific set.

For each replicate, the proportion of proxy alleles was calculated across ancient individuals, and the temporal association between proxy allele proportion and sample age was estimated using the same weighted Kendall's $\tau$ framework applied to the observed trait-specific markers. This generated a trait-specific null distribution of weighted Kendall's $\tau$ values expected under genome-wide demographic and background-selection effects. The observed weighted Kendall's $\tau$ was then compared with this null distribution using a one-sided Monte Carlo test according to the direction of the observed effect. For traits with positive observed $\tau$, empirical $P$ values were calculated as the proportion of null replicates with $\tau_{null} \geq \tau_{obs}$, whereas for traits with negative observed $\tau$, empirical $P$ values were calculated as the proportion of null replicates with $\tau_{null} \leq \tau_{obs}$.

Supplementary Fig. 28 shows the resulting null distributions for the six traits that remained significant after within-cluster FDR correction.

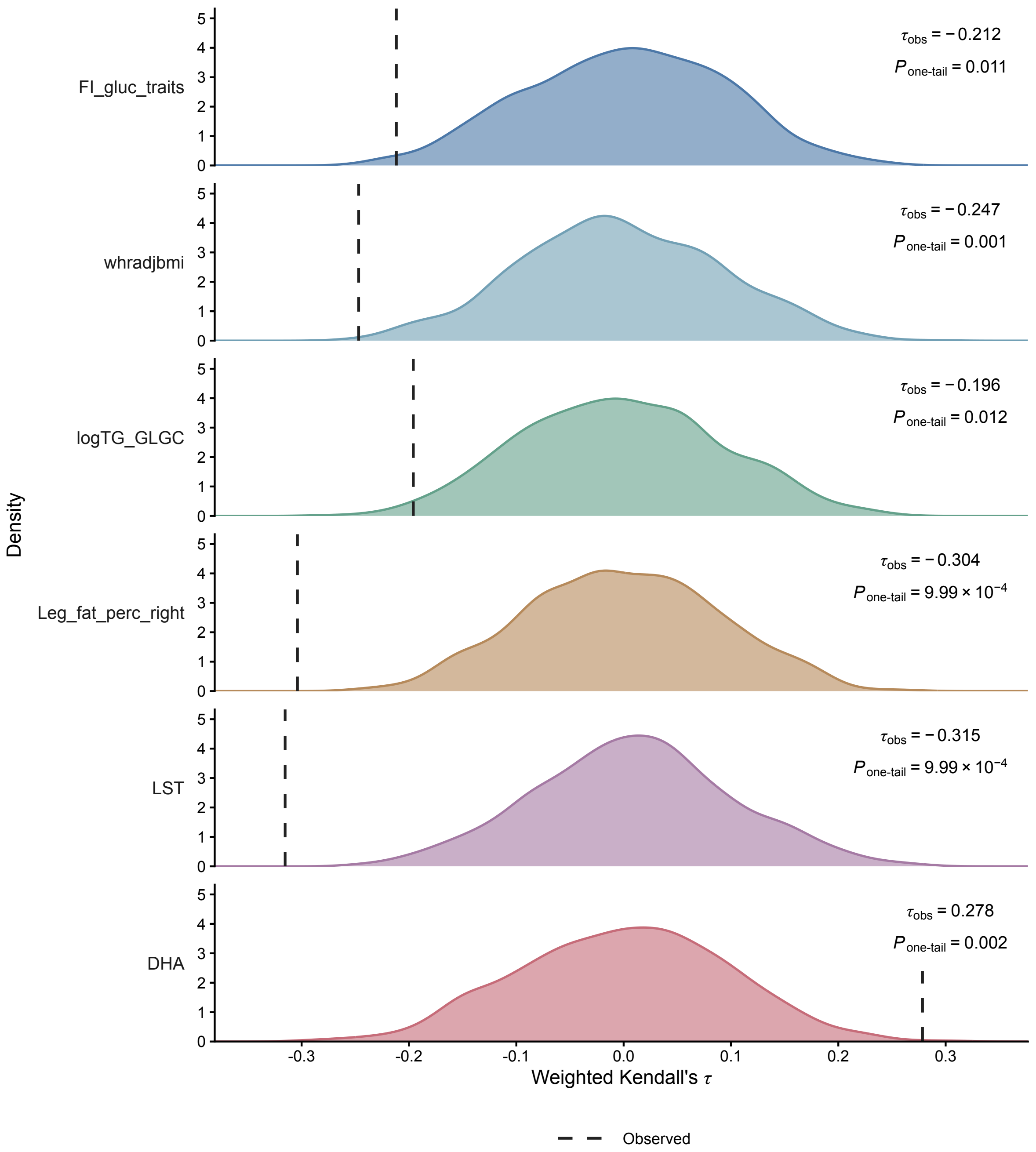


**Supplementary Fig. 28 | Empirical null distributions of temporal trajectories against genome-wide background variation.**

Empirical null distributions of weighted Kendall's $\tau$ generated from 1,000 matched proxy marker sets sampled from genome-wide variants linked to protein-coding genes, potentially sensitive to demographic history and background selection. Each distribution corresponds to one trait that remained significant after within-cluster FDR correction in the temporal analysis. Vertical dashed lines indicate the observed weighted Kendall's $\tau$ obtained from the corresponding trait-specific marker set. Empirical P values were calculated using a one-sided Monte Carlo test according to the direction of the observed effect. Traits shown are waist-to-hip ratio (WHR; $P = 0.001$), log-transformed triglycerides (logTG; $P = 0.012$), fasting insulin (FI; $P = 0.011$), right leg fat percentage ( $P = 9.99 \times 10^{-4}$), leisure screen time (LST; $P = 9.99 \times 10^{-4}$) and docosahexaenoic acid (DHA; $P = 0.002$).

#### Supplementary Note 7. Geographic partitioning of temporal trajectories

This section provides additional information on the geographic partitioning analysis used to evaluate whether pan-European temporal signals were driven by spatially heterogeneous trajectories. For each trait, the genetic algorithm searched across linear geographic barriers and identified the partition that maximized differentiation between the temporal patterns observed on either side of the barrier, while penalizing strongly unbalanced sample splits. This approach was designed to detect regional structure in trait-associated marker trajectories without imposing predefined geographic groupings.

A schematic overview of the algorithm is provided in Supplementary Fig. 29. Briefly, each candidate solution defined a linear barrier across Europe, which was used to assign ancient individuals to one of two geographic partitions. Regional differentiation was then evaluated from the fitted temporal trajectories of the two groups using the F-like statistic shown in the scheme. The GA was run with a population of 200 candidate solutions, a mutation probability of 0.2 and 100 generations. Parent selection followed a tournament strategy, in which five randomly sampled candidates competed for reproduction, with selection probabilities proportional to their fitness. Offspring were generated through recombination and mutation of the geographic barrier coordinates.

Supplementary Figs. 30 and 31 show representative regional temporal trajectories coloured by the fine genetic-cluster labels from Allentoft et al. Fasting insulin (FI) illustrates the Baltic–Black Sea-oriented partition observed for several traits, in which western groups were largely composed of WHG-related samples, whereas the north-eastern partition included Mesolithic populations with mixed EHG- and WHG-related ancestry. DHA illustrates a different west–east configuration, where hunter-gatherer-associated groups from both western and eastern Europe tended to depart from lower marker frequencies relative to later ancestry groups, followed by a stronger increase in the western partition.

**Algorithm 2: Genetic optimization of geographical regression splits**

**Input:** Dataset $D$ for one trait: longitude $x$, latitude $y$, age $t$ and risk-allele proportion $z$;
even population size $P$; mutation rate $\mu$; generations $G$; tournament size $k$; penalty strength $\lambda = 0.5$
**Output:** Best split $s_{\text{best}} = (x_0, y_0, x_1, y_1)$ and its fitness
**Function** `MapSplit`$(s, D)$:
- $b_h \leftarrow (y_1 - y_0)/(x_1 - x_0)$; $b_v \leftarrow (x_1 - x_0)/(y_1 - y_0)$
- **if** $|b_v| \le 0.1$ **then**
  - `// Near-vertical boundary: split by longitude`
  - $D_1 \leftarrow$ observations with longitude $< x_0$
- **else if** $|b_h| \le 0.1$ **then**
  - `// Near-horizontal boundary: split by latitude`
  - $D_1 \leftarrow$ observations with latitude $< y_0$
- **else**
  - `// Diagonal case: define the line through the two endpoints`
  - $a \leftarrow y_0 - b_h x_0$ `// Intercept`
  - $D_1 \leftarrow$ observations above the line latitude $= b_h$ longitude $+ a$
- **return** $(D_1, D_2)$, *where* $D_2$ *contains the remaining observations*

**Function** `Fitness`$(D_1, D_2)$:
- `// Compare risk-allele proportion over time between the two geographical groups`
- Fit $z_i = \alpha_r + \beta_r t_i + \varepsilon_i$ separately for $r = 1, 2$
- Obtain group size $n_r$, centered age sum of squares $SSx_r$, and residual sum of squares $SSE_r$
- $F \leftarrow \dfrac{(\beta_1 - \beta_2)^2}{\left(\frac{SSx_1 + SSx_2}{SSx_1 SSx_2}\right)\left(\frac{SSE_1 + SSE_2}{n_1 + n_2 - 4}\right)}$
- `// Symmetric penalty for group-size imbalance`
- **return** $f = F \exp\left(-\lambda \left|\log \dfrac{n_1}{n_2}\right|\right)$

`// Initialization: resample until both groups are non-empty`
$S \leftarrow \varnothing$
**for** $i \leftarrow 1$ **to** $P$ **do**
- **repeat**
  - Sample $x_0, x_1$ and $y_0, y_1$ independently from unique observed longitudes and latitudes, with replacement
  - $s_i \leftarrow (x_0, y_0, x_1, y_1)$; $(D_1, D_2) \leftarrow$ `MapSplit`$(s_i, D)$
- **until** *both* $D_1$ *and* $D_2$ *are non-empty*
- Append $s_i$ and its subsets to $S$

**foreach** $s_i \in S$ *in parallel* **do** $f_i \leftarrow$ `Fitness`$(D_{i1}, D_{i2})$
**for** $g \leftarrow 1$ **to** $G$ **do**
- `// Tournament reproduction`
- $S_{\text{new}} \leftarrow \varnothing$
- **for** $i \leftarrow 1$ **to** $P/2$ in parallel **do**
  - Sample a tournament $\mathcal{T}$ of $k$ population entries without replacement
  - $\pi_j \leftarrow \dfrac{f_j}{\sum_{\ell \in \mathcal{T}} f_\ell}$ for $j \in \mathcal{T}$
  - Select distinct parents $p_1, p_2$ using $\pi_j$, renormalizing after the first draw
  - `// Recombination weighted by parental fitness`
  - $w_1 \leftarrow f_{p_1}/(f_{p_1} + f_{p_2})$; $w_2 \leftarrow f_{p_2}/(f_{p_1} + f_{p_2})$
  - $c_1 \leftarrow w_1 s_{p_1} + w_2 s_{p_2}$; $c_2 \leftarrow w_2 s_{p_1} + w_1 s_{p_2}$
  - **foreach** *child* $c$ *among* $c_1, c_2$ **do**
    - `// Mutation: at most one coordinate; Gaussian SD is 5`
    - $B \sim \text{Bernoulli}(\mu)$
    - **if** $B = 1$ **then**
      - Select $\ell$ uniformly from $\{1, 2, 3, 4\}$; draw $X \sim \mathcal{N}(c[\ell], 5^2)$ and set $c[\ell] \leftarrow X$
    - Append $c$ and `MapSplit`$(c, D)$ to $S_{\text{new}}$
- **foreach** $c \in S_{\text{new}}$ *in parallel* **do** $f_c \leftarrow$ `Fitness`$(D_{c1}, D_{c2})$
- `// Elitist population update`
- $S \leftarrow$ the $P$ highest-fitness entries after concatenating $S$ and $S_{\text{new}}$

**return** *Highest stored fitness and corresponding split* $s_{\text{best}}$ *in* $S$

**Supplementary Fig. 29 | Genetic algorithm for geographic partitioning of temporal trajectories.**
Schematic overview of the genetic algorithm used to identify linear geographic barriers associated with differentiated temporal trajectories. Each candidate solution is represented by a vector $(x_0, y_0, x_1, y_1)$, defining the start and end coordinates of a linear barrier across Europe. The resulting slope and intercept are used to assign ancient individuals to two geographic partitions based on their longitude and latitude. For each partition, a linear model is fitted between the proportion of risk alleles and sample age, and the resulting group-level statistics are combined into the F-like statistic shown in the scheme. Candidate solutions with highly

imbalanced sample sizes are penalized using an exponential term that decreases the final fitness as the ratio between group sizes deviates from 1. Parent selection was performed using a tournament strategy, in which candidate solutions were randomly sampled according to `n_tournament` and allowed to compete for reproduction. Candidate fitness values were converted into sampling probabilities by dividing each fitness value by the sum of fitness values across candidates, so that higher-fitness solutions had a greater probability of being selected. This probability vector was then used to sample two parents with `rmultinom`. Offspring solutions were generated during recombination as a weighted average of the two parental barriers, with weights given by their fitness-derived sampling probabilities. Mutation sites were sampled from a binomial distribution with probability equal to the mutation rate, allowing stochastic perturbation of barrier coordinates across generations.

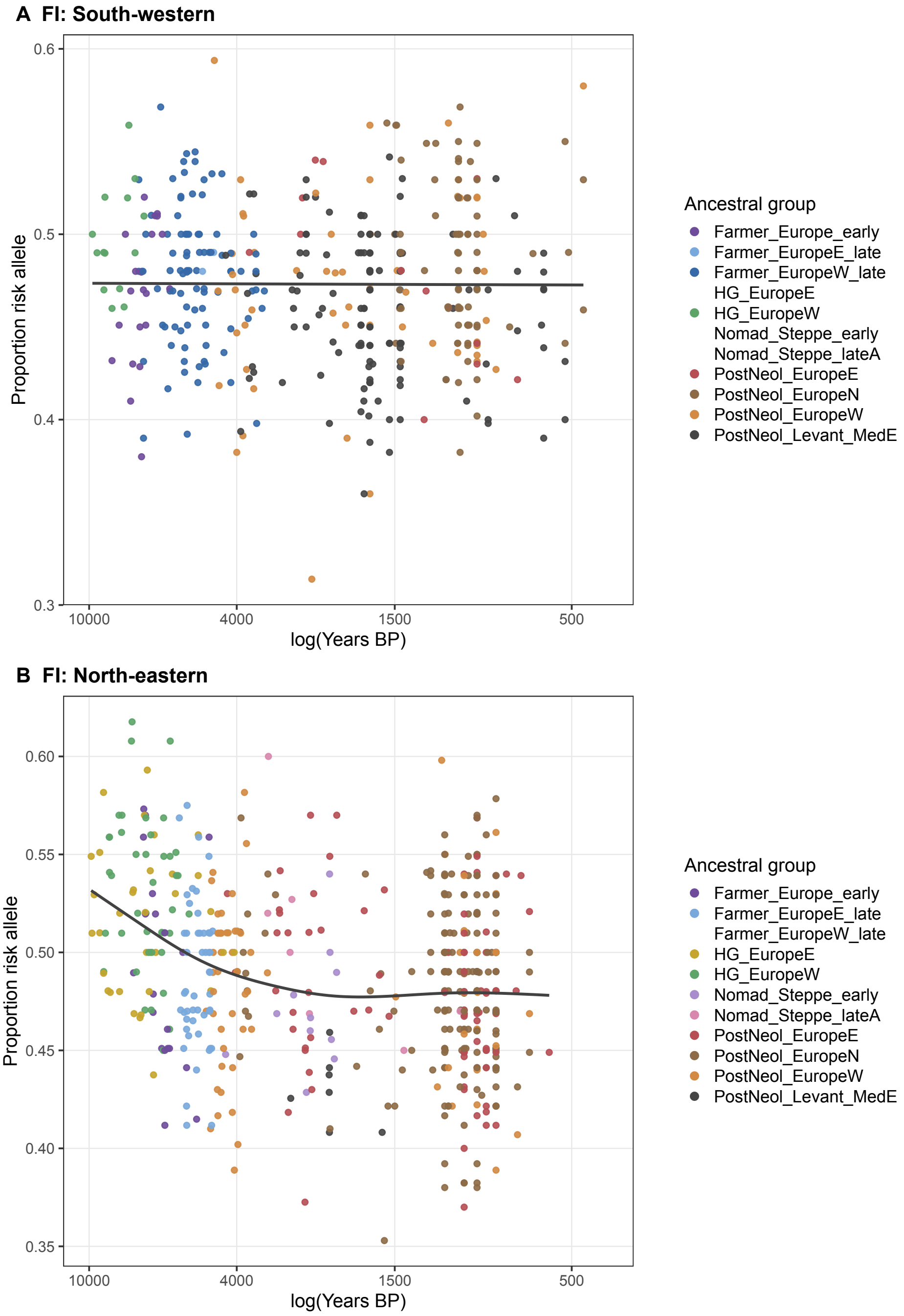


**Supplementary Fig. 30 | Regional fasting insulin trajectories under the Baltic–Black Sea partition.**

Temporal trajectories of fasting insulin (FI) marker load after partitioning ancient individuals according to the geographic barrier inferred by the genetic algorithm. **A**, south-western European partition; **B**, north-eastern European partition. Points represent ancient individuals coloured by the fine genetic-cluster labels from Allentoft et al. The x-axis represents sample age in years before present (BP) on a log scale, with labels shown in the original BP scale, and the y-axis represents the proportion of risk alleles. A GAM smooth is shown for visualization. This partition follows the Baltic–Black Sea axis and illustrates the stronger temporal trajectory observed in the north-eastern partition, which includes Mesolithic groups with mixed EHG- and WHG-related ancestry, compared with the predominantly WHG-related western partition.

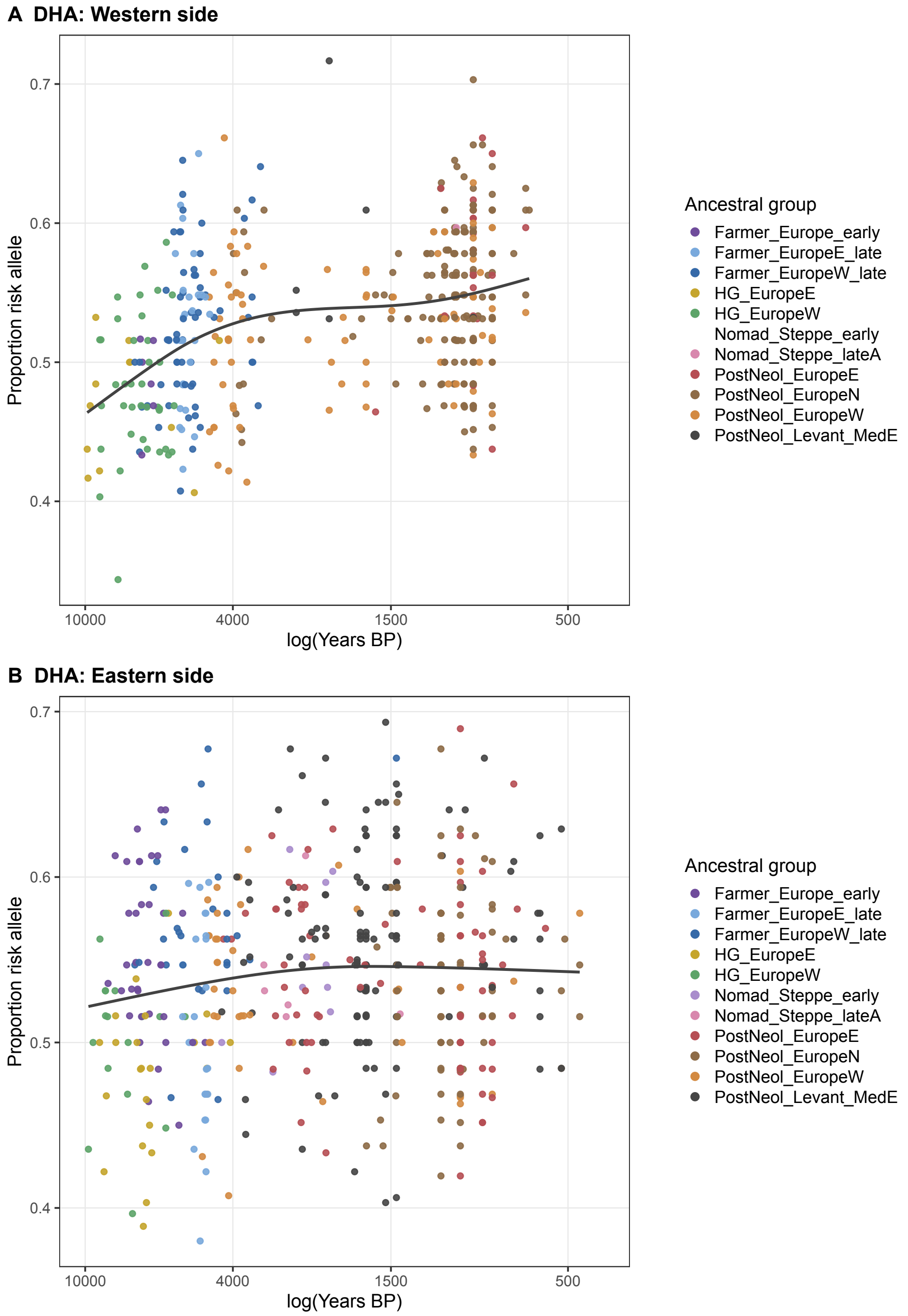


**Supplementary Fig. 31 | Regional DHA trajectories under the Gibraltar-Baltic partition.** Temporal trajectories of docosahexaenoic acid (DHA) marker load after partitioning ancient individuals according to the geographic barrier inferred by the genetic algorithm. **A**, western European partition; **B**, eastern European partition. Points represent ancient individuals coloured by the fine genetic-cluster labels from Allentoft et al. The x-axis represents sample age in years before present (BP) on a log scale, with labels shown in the original BP scale, and the y-axis represents the proportion of risk alleles. A GAM smooth is shown for visualization. This partition approximates a west–east barrier spanning the Gibraltar–Baltic axis and illustrates that hunter-gatherer-associated groups from both western and eastern Europe start from lower marker frequencies relative to later ancestry groups, with the strongest subsequent increase observed in the western partition.
